# ABA increases fatty acids levels in apple roots to boost colonization by arbuscular mycorrhizal fungi

**DOI:** 10.1101/2024.11.13.623435

**Authors:** Shan Jing, Mingjun Li, Lingcheng Zhu, Chunhui Li, Yuchao Li, Lijun Du, Xiaoyu Wei, Manrang Zhang, Baiquan Ma, Yongling Ruan, Fengwang Ma

**Affiliations:** State Key Laboratory for Crop Stress Resistance and High-Efficiency Production / Shaanxi Key Laboratory of Apple, College of Horticulture, Northwest A&F University, Yangling 712100, Shaanxi, China

**Keywords:** Abscisic acid, mycorrhizal symbiosis, ABF2, lipid synthesis and transfer, *Malus pumila* Mill

## Abstract

The roots of most land plants are in symbiosis with arbuscular mycorrhizal (AM) fungi. The fungus promotes nutrient uptake from the soil while receiving plant photosynthate as lipids and sugars. Nutrient exchange must be regulated by both partners, but the mechanisms underlying the regulation of lipid supplement from the plant to the AM fungus remain elusive. Here, we conducted a molecular study on the role of increased abscisic acid (ABA) levels during AM fungus infection in the roots of apple (*Malus* spp.). AM fungus induced the expression of two ABA synthesis genes, *MdNCED3.1* and *3.2*, in apple roots and increased the ABA content, which promoted the growth of the AM fungus. The effect of ABA on symbiosis was confirmed in transgenic apple roots either overexpressing or silencing *MdNCED3.1* or *MdNCED3.2*. Transcriptome analysis and transgenic manipulation revealed that the transcription factor MdABF2 played a key role in the ABA-mediated formation of symbiosis during AM infection and that MdABF2 could regulate the expression levels of genes related to fatty acid (FA) synthesis (e.g., *MdKASIII*) and translocation (such as *MdSTR2*) in apple roots. Activation of these genes boosted the levels of available fatty acids in the roots and increased the AM fungal colonization and arbuscule development in the roots. These results revealed a molecular pathway in which positive regulation of FA synthesis and transport by the ABA signaling pathway increases supplementation to AM fungi and promotes AM symbiosis.

## Introduction

Arbuscular mycorrhiza (AM) is a widespread symbiotic interaction formed between 80%–90% of land plant species and the *Glomeromycotina* fungi (Gadkar et al., 2001; Smith et al., 2011). From ecological and agronomic perspectives, this mutualistic symbiosis has garnered significant interest because AM enhances crop yield and quality by altering mineral nutrient uptake, stress resistance, and carbon or nutrient distribution in plants (Smith et al., 2011; Bennett and Groten, 2022; Shi et al., 2023; Wang et al., 2024). Typically, all mutualistic symbioses involve the exchange of nutrient “currencies” that confer adaptive advantages while potentially incurring costs. Therefore, the symbionts evolve regulatory mechanisms to prevent excessive exploitation by their trading partner (Kiers et al., 2011; Bennett and Groten, 2022). An optimal growth system requires a balanced transaction, and we seek to understand what the AM fungus helps the plant acquire and what costs does the plant incur to provide for the AM fungus.

The symbiotic relationship between AM fungi and host plants is built on mutual benefit, where the fungi provide nutrients to the plant while obtaining the carbon that is necessary for their growth in exchange. AM fungi require fixed carbon from plants as their food source, with plants relinquishing 4–20% of their photoassimilates, as carbohydrates and lipids, to the fungus (Jiang et al., 2017; Luginbuehl et al., 2017). In the past, detailed ^13^C labeling and nuclear magnetic resonance tracing studies suggested that hexoses are the primary forms of carbon transferred from plants to fungi (Shachar-Hill et al., 1995). It was believed that plants only provided sugars to fungi, and that the AM fungi utilized these sugars as precursors for lipid biosynthesis (Pfeffer et al., 1999). However, *de novo* fungal fatty acid (FA) synthesis is only observed in colonized roots, not in extraradical hyphae or spores (Trépanier et al., 2005). In recent years, significant progress has been made in understanding this metabolism. Jiang et al. (2017), using techniques like isotope labeling and split-root systems, demonstrated that FAs are the primary carbon supplied from plants to AM fungi. The AM fungi primarily store carbon as triacylglycerol (TAG), and most FAs in AM fungi are composed of 16:0 (palmitic acid) 16:1 (palmitoleic acid) species in the mycorrhized roots (Trépanier et al., 2005; Keymer et al., 2017). After penetrating the plant roots, AM fungi could induce FA synthesis and transport to meet its own food requirements (Jiang et al., 2017; Luginbuehl et al., 2017). In plant root cells, the synthesis of lipids starts with acetyl-CoA (Shin and Ohlrogge, 2009). Sixteen-carbon FA chains can be synthesized through the ketoacyl-ACP synthase (KAS) and FA synthase system (FAS) and released from FAS by acyl-ACP thioesterases (FatM). The saturated 16:0 FA is converted into 2-monoacylglycerol (2-MAG) molecules by RAM2 within the colonized plant cell and transported into the symbiotic space by the STR/STR2-mediated lipid export pathway (Jiang et al., 2017). The lipid is further taken up by unknown lipid transporters in AM fungi. However, it is unknown how to activate lipid synthesis and translocation in plant roots after fungal infection.

As part of the symbiosis, the AM fungus penetrates the plant roots and forms structures called hyphae and arbuscule within the plant cells. AM symbiosis causes changes in the levels of thousands of transcripts and metabolites in plants (Bravo et al., 2016; Jing et al., 2022; Shao et al., 2023). In addition, plant hormones, as important regulators of growth and development as well as response to environmental conditions, have also been shown to play key roles in regulating the interaction between plants and AM fungi (Liu et al., 2020; Jing et al., 2022). Almost every known plant hormone contribute to AM formation and function, from early symbiosis pre-signaling to later stages, including the morphological root cell changes necessary for the acceptance of fungal cells (Küster and Gutjahr, 2017; Pozo et al., 2020). The germination of fungal spores and hyphal growth and root colonization are initiated by gibberellins (GAs) and auxins (Liao et al., 2018). On the other hand, salicylates and ethylene play negative roles during fungal invasion and root colonization (Blilou et al., 2000; Rodolfo et al., 2009). Interestingly, ABA, a growth-inhibiting hormone, has a positive impact on AM formation and function. ABA levels are drastically induced by AM fungi (Jing et al., 2022), suggesting that ABA might play a crucial role in AM symbiosis. Studies have reported that ABA facilitates AM fungal growth and promotes arbuscule formation and hyphal elongation (Herrera-Medina et al., 2007; Charpentier et al., 2014;Liu et al., 2016). Numerous studies have shown that ABA promotes AM fungal symbiosis with crops (Charpentier et al. 2014; Miransari et al. 2014; Han et al. 2023), but the pathway through which AM fungi actually induce ABA to promote colonization is unclear.

In this study, we investigated the role of increased ABA levels in regulating lipid supplementation from apple plants to arbuscular mycorrhizal fungus *Rhizophagus irregularis*. It was observed that increased ABA concentration in apple root promotes AM fungal growth. Inoculation with the AM fungus induced the expression of *MdNCED3.1*/*3.2*, thereby increasing ABA synthesis in apple roots, which regulates FA synthesis and transport via transcript factor MdABF2, thereby boosting the supplementation of FA and enhancing the symbiotic efficiency of AM. These results revealed a novel molecular mechanism by which ABA promotes AM symbiosis. The ABA signaling pathway positively regulates FA synthesis and transport, thereby increasing carbon supplementation to AM fungi and promoting AM symbiosis.

## Results

### During AM symbiosis, the ABA content increases in apple roots

As shown in our previous report (Jing et al., 2022), AM symbiosis induces obvious changes in phytohormone-related transcripts and metabolites in the roots of apple seedlings (Figure 1B and 1C). Among the plant hormones, ABA content increased approximately 10-fold after inoculation with mycorrhizae (Figure 1C). The expression of genes (especially *MdNCED*) related to ABA synthesis also followed increasing trends, with significant up-regulation after inoculation (Figure 1D). The transcript levels of *MdNCED3.1* and *MdNCED3.2* were much higher than those of other members of the NCED family (Supplemental Figure 1 and Supplemental Figure 2). These results suggested that mycorrhizal symbiosis could induce an increase in ABA synthesis-related transcription in apple root.

**Figure 1.**
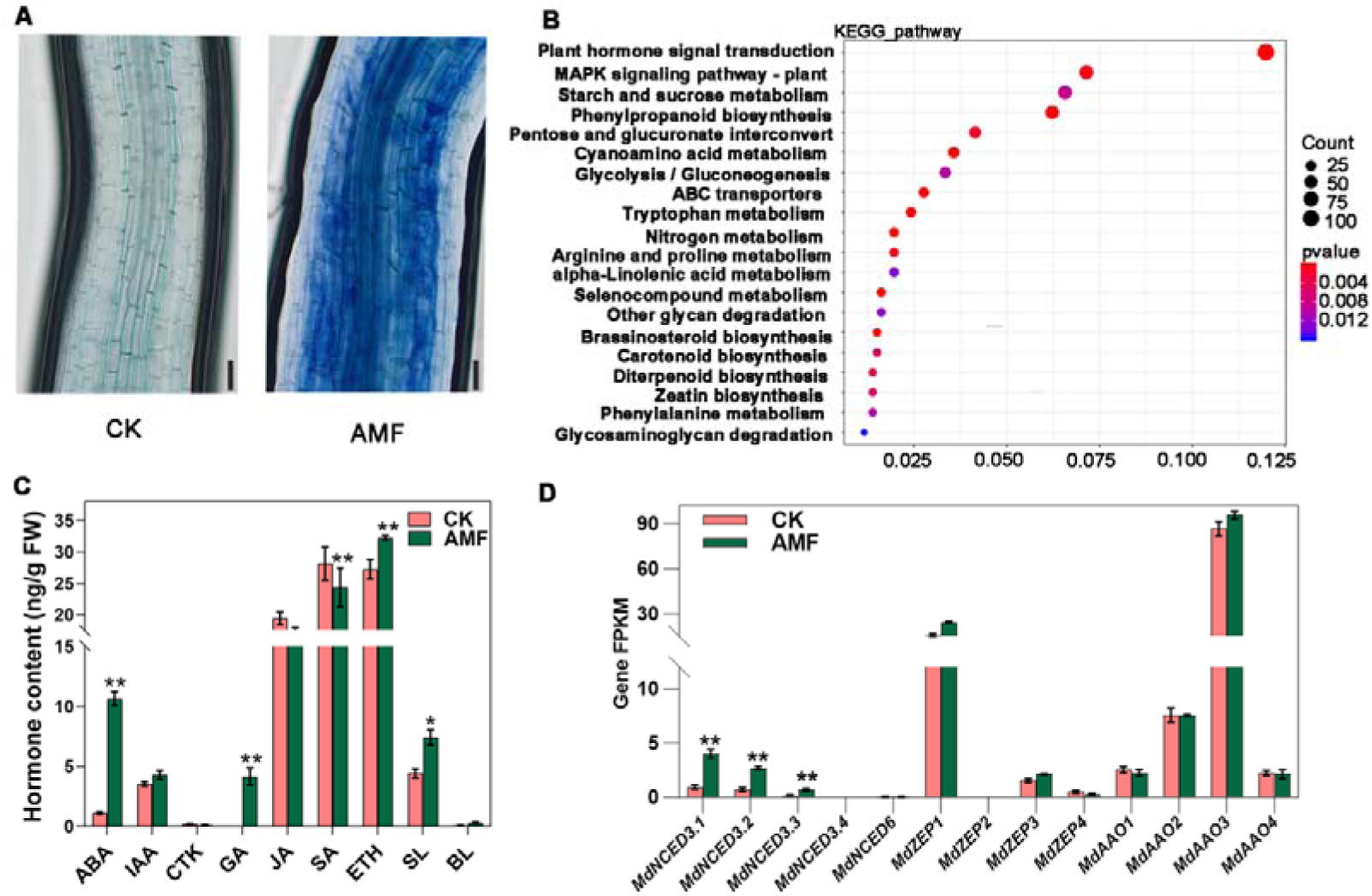
Roots of M26 (*Malus pumila* Mill.) plants, uninoculated or inoculated with arbuscular mycorrhizae (AM) fungus and analyzed after 60 days via transcriptomics and metabolomics. **(A)** Staining of mycorrhizae in uninoculated (CK) and inoculated (AMF) M26 roots. Scale bars, 100 µm. **(B)** KEGG-based enrichment analysis of differentially expressed genes in the root transcriptome. **(C)** Hormone content in M26 plants with and without inoculation. **(D)** Expression levels of ABA synthesis-related genes, as measured by FPKM from RNA-seq, in both uninoculated and inoculated M26 plants. The bars represent the mean value ± SD (n = 3 independent biological replicates). Mixed samples from three M26 plants were as one replicate. The asterisks indicate significant differences as assessed by one-way ANOVA (two-sided Student’s *t*-test; \*\**P* < 0.01, \**P* < 0.05).

### ABA helps the colonization of AM fungi in apple root

To determine if ABA has a role in symbiosis between apple roots and AM fungus, we first investigated the influence of exogenous ABA and ABA inhibitors on colonization by AM fungus *Rhizophagus irregularis*. Thirty days after inoculation, 50 µmol/L ABA or 50 µmol/L fluridone (Flu, an ABA synthesis inhibitor) was added to the water used on the apple roots. After 15 days of treatment, there was a notable increase in the lateral roots of plants treated with ABA, whereas plants treated with Flu exhibited a significant decrease in lateral roots (Fig. 2a, b, d, f, and g). ABA treatment significantly promoted AM symbiosis, whereas the Flu treatment significantly inhibited AM symbiosis, especially the development of arbuscules and growth of inter/intracellular hyphae (Fig. 2c, h). These findings indicated that ABA is an important part of AM symbiosis.

**Figure 2.**
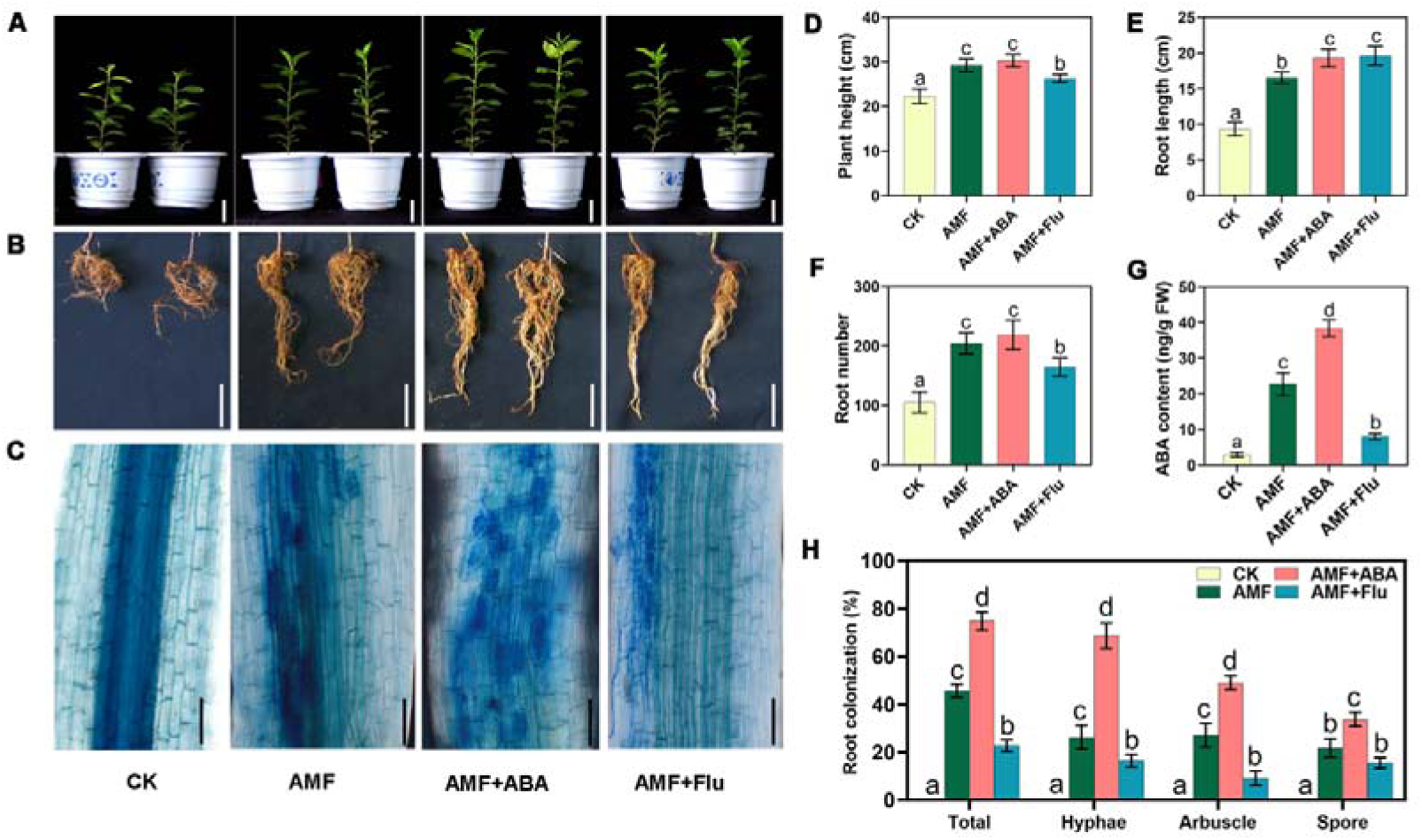
Exogenous ABA positively regulates AM symbiosis in M26 apple (*Malus pumila* Mill.) plants. **(A–C)** Phenotypes of 45-day-old M26 plants under control (CK), arbuscular mycorrhizae-inoculated (AMF), inoculated plus ABA-treated (AMF+ABA), and inoculated plus ABA inhibitor-treated (AMF+Flu) conditions. **(A)** Plant height. Scale bars, 5 cm. **(B)** Root structure. Scale bars, 5 cm. **(C)** Trypan blue staining of the fungus to reveal arbuscule morphology. Scale bars, 100 µm. **(D)** Height of M26 plants. **(E)** Root length of M26 plants. **(F)** Root number of M26 plants. **(G)** ABA content in roots of M26 plants. The bars represent the mean value ± SD (n = 3 independent biological replicates). Mixed samples from three M26 plants were as one replicate. **(H)** Quantification of mycorrhizal colonization levels. FW, fresh weight; CK, Uninoculated plants without ABA treatment for 45 days, as control. AMF, *Rhizophagus irregularis*-inoculated plants without ABA treatment for 45 days. AMF+ABA, *Rhizophagus irregularis*-inoculated plants were sprayed with exogenous ABA (50 µmol/L) for 15 days after inoculation for 30 days. AMF+ Flu, *Rhizophagus irregularis*-inoculated plants were sprayed with exogenous Flu (50 µmol/L ABA synthesis inhibitor fluoridone) for 15 days after inoculation for 30 days. (**D**, **E**, **F**, and **H)** The bars represent the mean value ± SD (n = 9 independent biological replicates). (**D**, **E**, **F**, **G**, and **H)** Different letters indicate significant difference (analysis of variance [ANOVA]), Duncan’s multiple range test; *P* < 0.05).

The ABA synthesis genes *MdNCED3.1* and *MdNCED3.2* were both upregulated in apple roots by AM fungus inoculation (Figure 1D and Supplemental Figure 2). We created independent *MdNCED3.1* and *MdNCED3.2* overexpression and silencing hairy root lines of the common seedling (*Malus hupehensis* Rhed, which is a triploid, typical apomixis-type of *Malus* species) Overexpression of *MdNCED3.1* or *MdNCED3.2* in apple roots increased ABA content (Supplemental Figure 3D, 3E, and 3I; Supplemental Figure 4), and significantly promoted AM symbiosis after mycorrhizal inoculation in apple roots (Figure 3C and 3I). In contrast, interference of *MdNCED3.1* or *MdNCED3.2* inhibited the increase of ABA content after fungal inoculation in the roots, reduced the infection rate by 50% compared to wild type roots, and significantly reduced AM symbiosis, especially the formation of arbuscules (Figure 3C and 3I). And, after adding exogenous ABA to the *RNAi-MdNCED3.1/3.2* lines, the arbuscular phenotype mostly recovered (Figure 3D and 3J). These results suggest that AM fungi induce *MdNCED3.1/3.2* expression to promote endogenous ABA synthesis in apple roots, which is necessary for efficient colonization by AM fungi.

**Figure 3.**
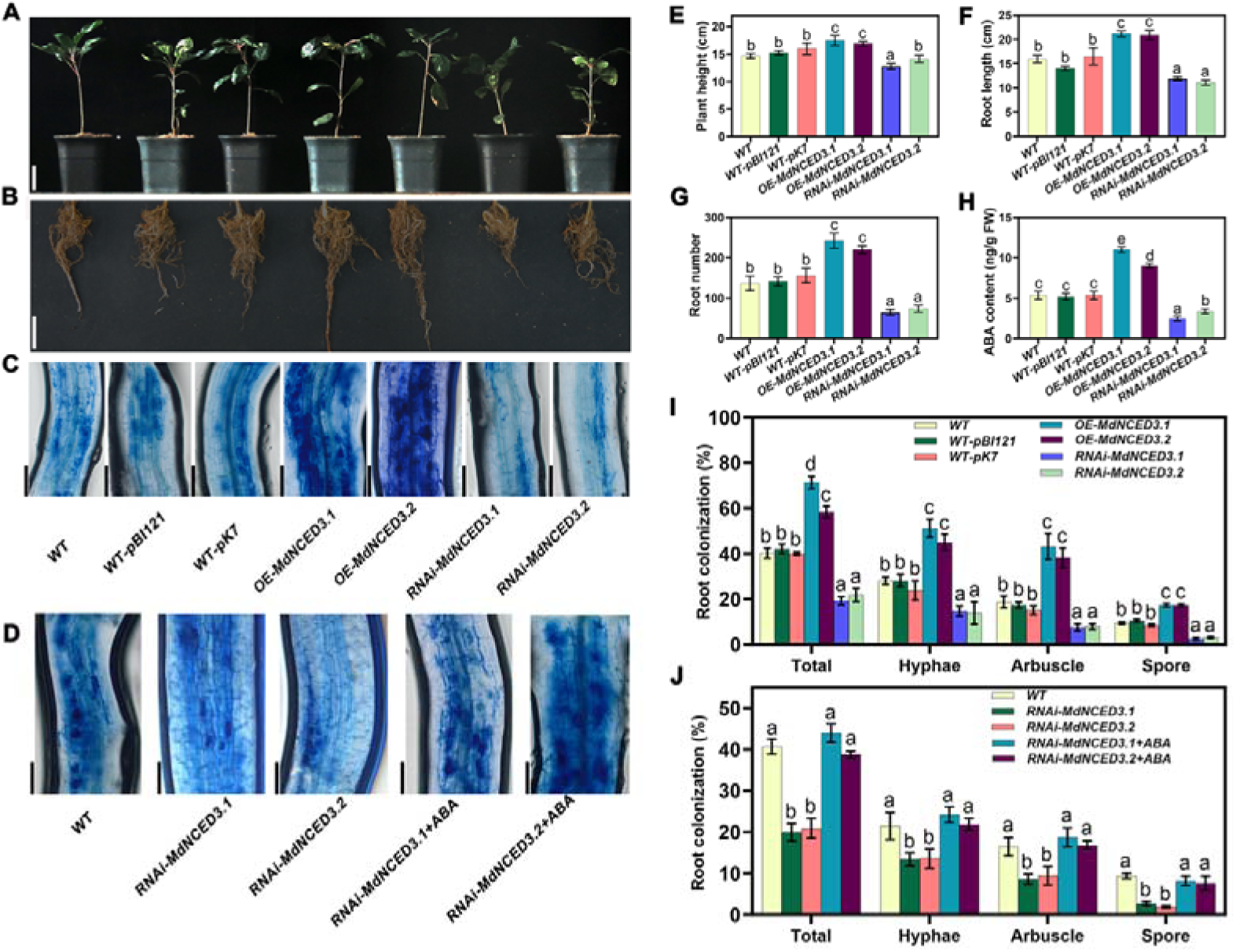
*MdNCED3.1/3.2* positively regulates AM symbiosis in apple (*Malus hupehensis* Rhed) seedlings. **(A–C)** Above-ground growth phenotypes **(A)**, root growth phenotypes **(B)**, and trypan blue staining of the fungus to reveal the arbuscule morphology **(C)** of 60-day-old WT seedlings and hairy roots transformed with WT-pBI121(empty vector), WT-pK7, OE-*MdNCED3.1*, OE-*MdNCED3.2*, RNAi-*MdNCED3.1*, or RNAi-*MdNCED3.2*. **(D)** Trypan blue staining of the fungus to reveal the arbuscule morphology in 75-day-old WT, RNAi-*MdNCED3.1*, and RNAi-*MdNCED3.2* roots treated with ABA. **(E)** Height of apple seedlings carrying transgenic hairy roots after AM spore inoculation. **(F)** Root length of apple seedlings carrying transgenic hairy roots after AM inoculation. **(G)** Root number of apple seedlings carrying transgenic hairy roots after AM inoculation. **(H)** ABA content in the root of apple seedlings carrying transgenic hairy roots after AM inoculation. The bars represent the mean value ± SD (n = 3 independent biological replicates). Mixed samples from three apple seedlings carrying transgenic hairy roots were as one replicate. **(I)** Quantification of mycorrhizal colonization levels in apple transgenic roots. **(J)** Quantification of mycorrhizal colonization levels of apple transgenic roots treated with ABA. RNAi-*MdNCED3.2*+ABA indicates *MdNCED3.2*-RNA interference roots that were sprayed with exogenous ABA (50 µmol/L) for 15 days beginning 45 days after fungal spore inoculation. **(A)** Scale bars, 5 cm. **(B)** Scale bars, 5 cm. **(C), (D)** Scale bars, 100 µm. FW, fresh weight. The genetic modification of the apple seedlings was only applied to the root. (**E**, **F**, **G**, **I** and **J)** The bars represent the mean value ± SD (n = 9 independent biological replicates). (**E**, **F**, **G**, **H**, **I** and **J)** Different letters indicate significant difference (analysis of variance [ANOVA]), Duncan’s multiple range test; *P* < 0.05).

### The ABA signal promotes AM symbiosis via the transcription factor MdABF2

ABA mainly functions via regulating expression of target genes, including transcription factors that further the signaling pathways (Yang et al., 2023). Analysis of our transcriptomic data of genes in the ABA signal transduction pathway revealed that a transcription factor, *MdABF2*, was significantly up-regulated in roots inoculated with AM fungus compared with uninoculated roots (Supplemental Figure 5). The expression level of *MdABF2* was also significantly increased in the roots of exogenously treated with ABA and in the *MdNCED3.1/3.2*-overexpression lines (Supplemental Figure 6).

To confirm whether ABF2 is an intermediary between ABA and AM symbiosis, we generated transgenic roots overexpressing and silencing *MdABF2* via the hairy root *Agrobacterium* transformation (Supplemental Figure 7). Sixty days after AM inoculation, the *MdABF2*-overexpression lines showed better root growth and aboveground development (Figure 4A, 4B, 4D, 4E, and 4F) and greater AM symbiosis, especially in regards to the number of arbuscules (Figure 4C and 4H), than WT. In contrast, *MdABF2-*RNAi lines had fewer arbuscules (Figure 4C and 4H). These results suggest that MdABF2 plays a key role in ABA-mediated progression of AM symbiosis.

**Figure 4.**
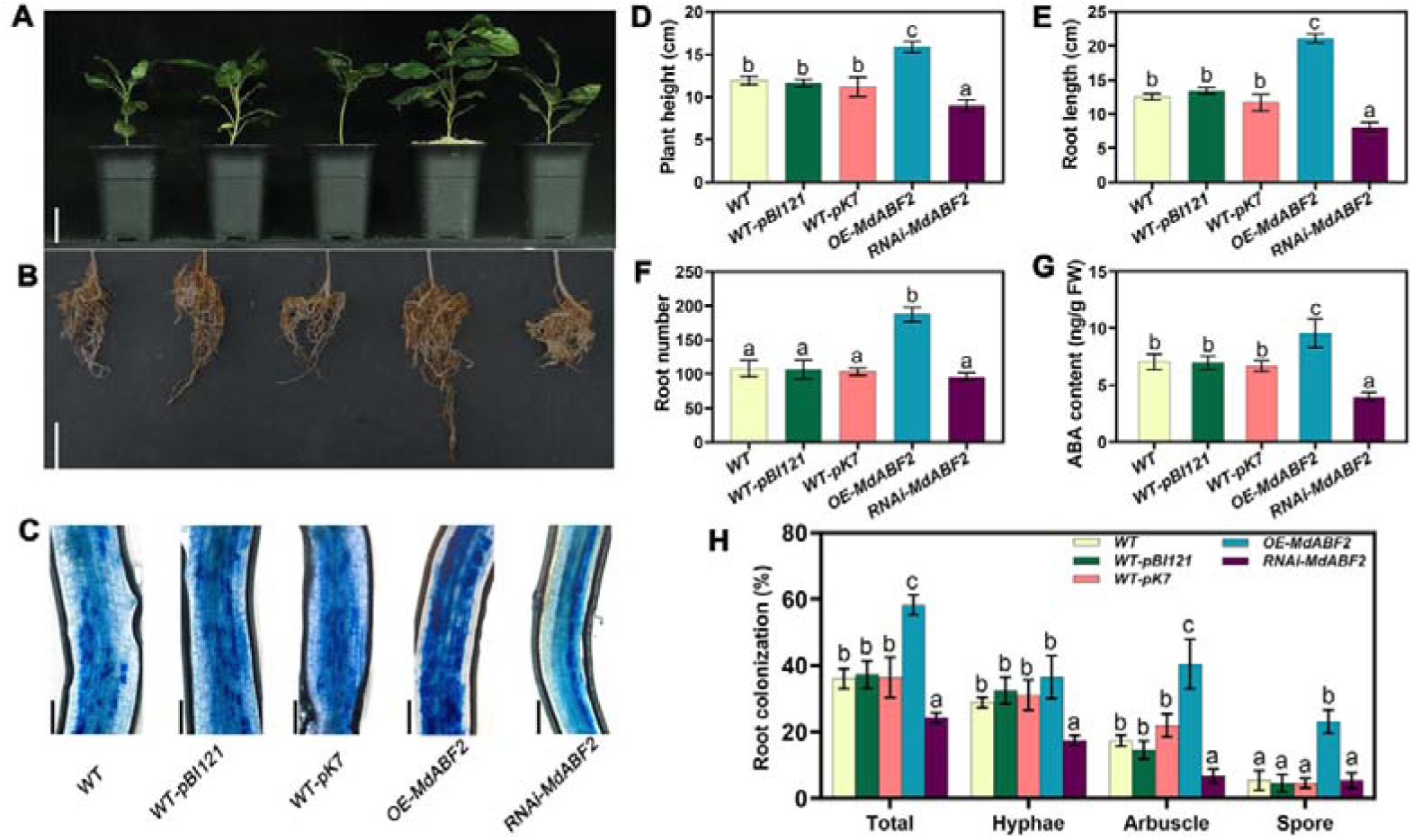
MdABF2 positively regulates AM symbiosis in apple (*Malus hupehensis* Rhed) hairy roots. **(A–C)** Above-ground growth phenotypes **(A)**, root growth phenotypes **(B)**, and trypan blue staining of the fungus to reveal arbuscular morphology **(C)** of 60-day-old WT, WT-pBI121 root transgenic lines, WT-pK7 transgenic lines, OE-*MdABF2* transgenic lines, and RNAi-*MdABF2* transgenic lines of apple rootstock. Indices were determined in five apple lines: **(D)** Height. **(E)** Root length. **(F)** Root number. **(G)** ABA content in the root. The bars represent the mean value ± SD (n = 3 independent biological replicates). Mixed samples from three apple seedlings carrying transgenic hairy roots were as one replicate. **(H)** Quantification of mycorrhizal colonization levels. **(A)** Scale bars, 5 cm. **(B)** Scale bars, 5 cm. **(C)** Scale bars, 100 µm. FW, fresh weight. The genetic modification of the apple seedlings was only applied to the root. The bars represent the mean value ± SD. (**D**, **E**, **F**, and **H)** The bars represent the mean value ± SD (n = 9 independent biological replicates). (**D**, **E**, **F**, **G**, and **H)** Different letters indicate significant difference (analysis of variance [ANOVA]), Duncan’s multiple range test; *P* < 0.05).

### ABA signal influences FA levels during AM symbiosis

Fatty acids are the primary carbon supplied from plant roots to AM fungus, and their levels are one key to development of the symbiosis (Jiang et al., 2017; Rich et al., 2017; Ivanov and Harrison, 2024). Therefore, we measured the FA and TAG contents in the roots of wild-type M26 with exogenous ABA and Flu treatments and in the various transgenic hairy root lines. The results indicated that increased ABA content or signaling increased the FA and TAG contents in the roots, and disruption in ABA synthesis or signaling decreased the FA and TAG contents (Figure 5A and 5B). And changes in MdNCED expression level are linked with FA and TAG content (Figure 5C and 5D). While changes in *MdABF2* expression level also correlated with changes in FA and TAG content (Figure 5E and 5F).

**Figure 5.**
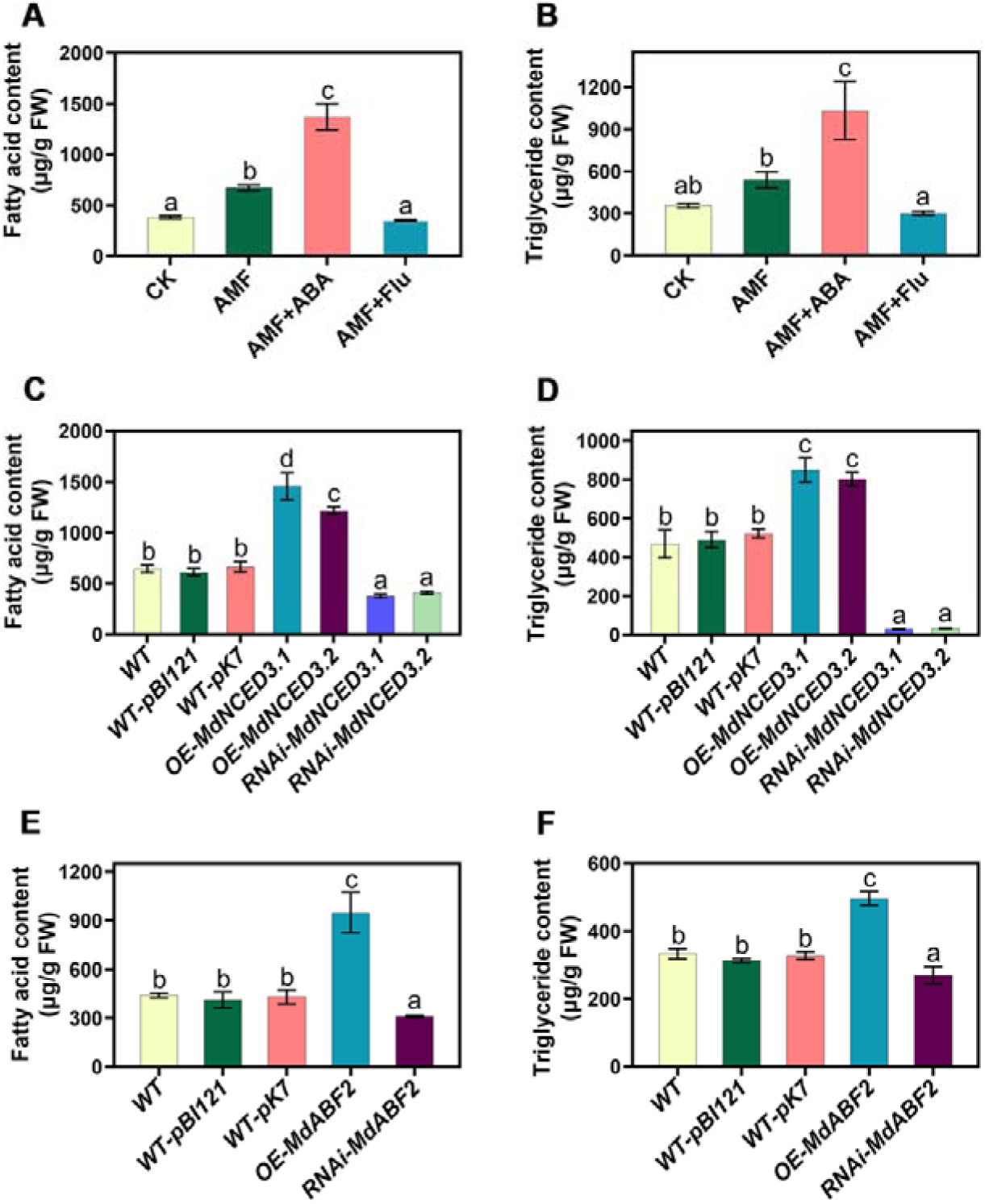
Fatty acid and triglyceride contents in apple roots following *R*. *irregularis* infection. **(A and B)** Non-transgenic M26 apple rootstock roots were tested for fatty acid **(A)** and triglyceride **(B)** content under CK, AMF, AMF+ABA, and AMF+Flu conditions. **(C), (D)** Sixty-day-old apple (*Malus hupehensis* Rhed) roots were tested for fatty acid **(C)** and triglyceride **(D)** content in WT and the transgenic lines WT-pBI121, WT-pK7, OE-*MdNCED3.1*, *OE-MdNCED3.2*, RNAi-*MdNCED3.1* and *RNAi-MdNCED3.2*. **(E), (F)** For the transcription factor lines, fatty acid **(E)** and triglyceride **(F)** content were determined in WT, WT-pBI121, WT-pK7, *OE-MdABF2*, and *RNAi-MdABF2*. FW, fresh weight. CK, Non-mycorrhizal inoculated plants without ABA treatment for 45 days as a control. AMF, Mycorrhizal-inoculated plants without ABA treatment for 45 days. AMF+ABA, Mycorrhizal-inoculated plants were prayed with exogenous ABA (50 µmol/L) for 15 days after inoculation for 30 days. AMF+ Flu, mycorrhizal-inoculated plants were prayed with exogenous Flu (50 µmol/L ABA synthesis inhibitor fluoridone) for 15 days after inoculation for 30 days. The bars represent the mean value ± SD (n = 3 independent biological replicates). (**A** and **B**) Mixed samples from three M26 plants were as one replicate. (C, D, E, and F) Mixed samples from three apple seedlings carrying transgenic hairy roots were as one replicate. Different letters indicate significant difference (analysis of variance [ANOVA]), Duncan’s multiple range test; *P* < 0.05).

### The transcription factor MdABF2 regulated expression of genes related to FA synthesis in apple root

We next analyzed the differentially expressed genes (DEGs) related to lipid metabolism based on two transcriptome databases, one of which compared the roots of mycorrhizal and non-mycorrhizal plants, and the other of which was from mycorrhizal roots of transgenic lines overexpressing or silencing *MdABF2*. The shared DEGs from the two transcriptomes included *MdKASI*, *MdKASI-1*, *MdKASIII*, *MdRAM2*, *MdRAM2-1*, *MdSTR2*, and *MdWRI3*. These genes were significantly upregulated after inoculating roots with AM fungus, and this upregulation required the MdABF2 transcription factor (Supplemental Figure 9; Supplemental Data 2). The promoter regions of these seven genes each contained at least one ABRE element, which is a potential binding domain for ABF (Supplemental Figure 10A). And *MdSTR2* promoter regions contained AW-box, which is a potential binding domain of WRI3 (Supplemental Figure 10C).

Preliminary yeast one-hybrid (Y1H) assays showed that MdABF2 can activate the FA synthesis genes *MdKASI*, *MdKASI-1*, *MdKASIII*, *MdRAM2*, and *MdRAM2-1*, the translocation protein gene *MdSTR2*, and the transcriptional factor *MdWRI3* (Supplemental Figure 10B). Likewise, MdWRI3 can activate *MdSTR2* (Supplemental Figure 10D). The genes *MdKASIII*, *MdWRI3*, and *MdSTR2* showed higher transcript abundance in apple roots, and much higher upregulation with AM inoculation (Supplemental Figure 9 and Supplemental Figure 11).

To further verify whether MdABF2 can regulate the expression of genes related to FA synthesis, translocation, and transcriptional regulation, or MdWRI3 can regulate the expression of *MdSTR2* through physical binding to their promoters, GUS-promoter fusion, dual luciferase, and electromobility shift assays were conducted between MdABF2 and *MdKASIII*, *MdWRI3*, and *MdSTR2* promoter sequences and between MdWRI3 and *MdSTR2* promoter sequences. GUS enzyme activity assays showed that co-expression of *MdSTR2-Pro:GUS* with *35S:MdWRI3* and of either *MdWRI3-*, *MdKASIII-*, or *MdSTR2-Pro:GUS* with *35S:MdABF2* in tobacco leaves significantly increased GUS staining (Figure 6A). Luciferase assays confirmed that co-expression of the transcription factor *35S:MdABF2* significantly increased luciferase activity when the *LUC* gene was driven by the promoter of either *MdWRI3*, *MdKASIII*, or *MdSTR2* in tobacco leaves. Likewise, co-expression of the transcription factor *35S:MdWRI3* and *MdSTR2-Pro:LUC* significantly increased luciferase activity (Figure 6B and Supplemental Figure 12). *In vitro*, EMSA experiments showed that the *MdKASIII*, *MdWRI3*, and *MdSTR2* promoter fragments containing ABREs were bound by recombinant MdABF2-His and resulted in a mobility shift. Similar mobility shifts were observed for interactions between MdWRI3-His protein and the *MdSTR2* promoter fragment (Figure 6C). Overall, these results indicated that MdABF2 can directly bind to the promoter sequences of *MdKASIII, MdWRI3*, and *MdSTR2* and that MdWRI3 can directly bind to the promoter sequence of *MdSTR2*, thereby promoting transcription of FA synthesis and transport genes.

**Figure 6.**
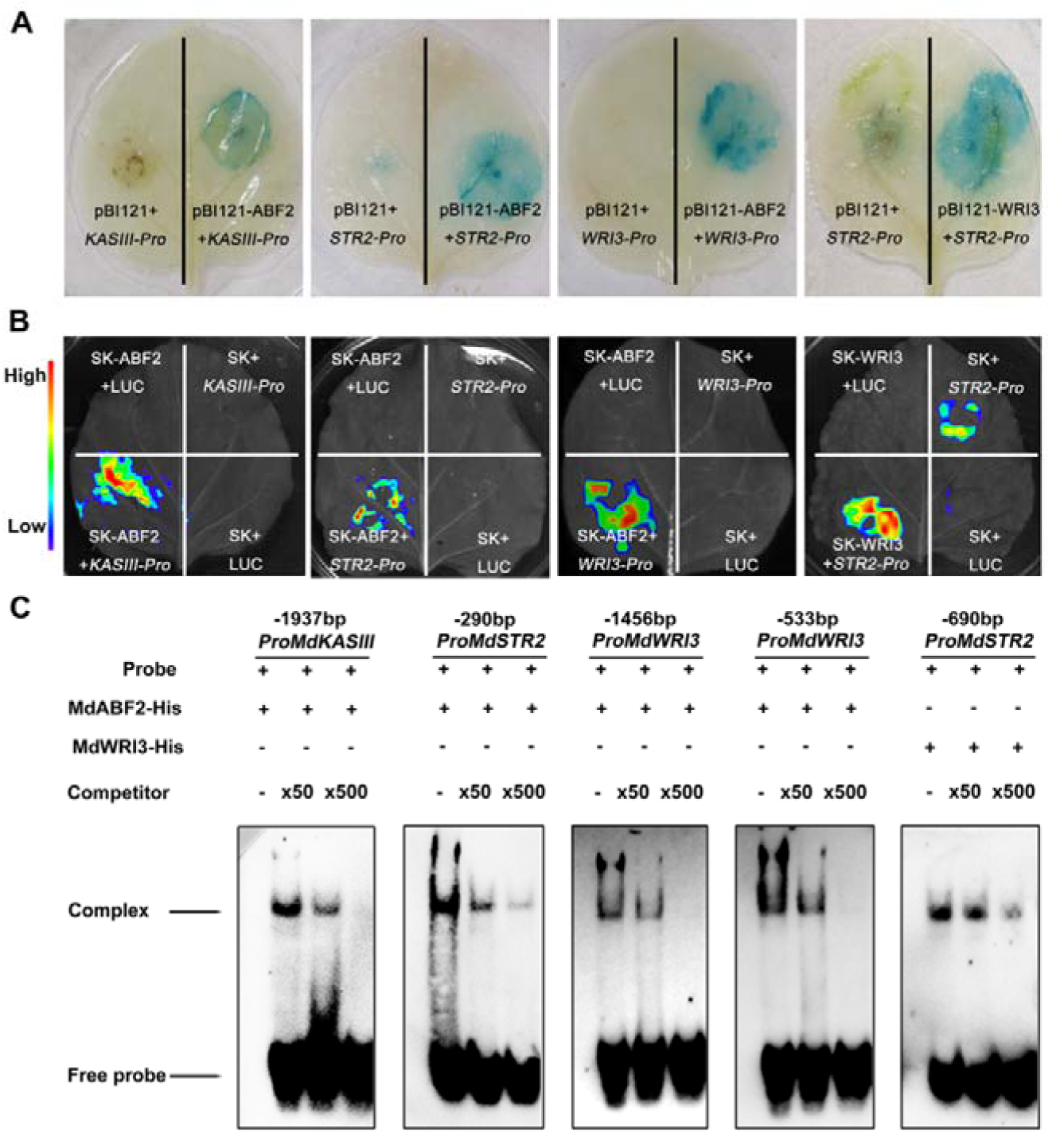
MdABF2 regulates the expression of *MdKASIII*, *MdSTR2*, and *MdWRI3*, and MdWRI3 regulates the expression of *MdSTR2*. **(A)** GUS staining reveals the interaction between MdABF2 and the promoters of *MdKASIII*, *MdSTR2*, and *MdWRI3*, as well as the interaction between MdWRI3 and the promoter of *MdSTR2*. Promoter-GUS fusions (*MdKASIII-Pro:GUS*, *MdWRI3-Pro:GUS*, or *MdSTR2-Pro:GUS*) and either the empty pBI121 vector or the pBI121 vector carrying the transcription factor *MdABF2* gene were transiently expressed in tobacco leaves. *MdSTR2-Pro:GUS* was co-transformed with the *MdWRI3* overexpression vector (pBI121-Pro35S:MdWRI3) or an empty vector control (pBI121). **(B)** LUC assays in tobacco leaves reveal the interaction between MdABF2 and the promoters of *MdKASIII*, *MdSTR2*, and *MdWRI3*, as well as the interaction between MdWRI3 and the promoter of *MdSTR2*. Promoter-LUC fusions (*MdKASIII-Pro:LUC*, *MdWRI-Pro3:LUC*, *MdSTR2-Pro:LUC*, and empty vector (pGreenII 0800-LUC) were co-transformed with either the MdABF2 overexpression vector (*Pro35S:MdABF2*) or an empty vector control (pGreenII 62-SK). *MdSTR2-Pro: LUC* or the empty vector (pGreenII 0800-LUC) were co-transformed with either the MdWRI3 overexpression vector (*Pro35S:MdWRI3*) or an empty vector control (pGreenII 62-SK). **(C)** EMSAs reveal interactions between MdABF2 or MdWRI3 and 5’-biotin-labeled probes of portions of the *MdKASIII*, *MdSTR2*, and *MdWRI3* promoters. Competing unlabeled probes without a biotin label were used at two concentrations.

### Synthesis and transport of FAs supports AM symbiosis in apple root

To determine whether *MdKASIII* affects FA synthesis and AM symbiosis in apple roots, we generated transgenic hairy roots either overexpressing or silencing *MdKASIII*. Overexpression of *MdKASIII* increased the contents of FA and TAG, and promoted AM symbiosis in apple roots, particularly arbuscule formation. Silencing of *MdKASIII* reduced fatty acid and TAG contents and AM colonization (Figure 7). The results indicated that KASIII affects AM symbiosis by altering FA accumulation. Likewise, overexpression of the transporter *MdSTR2* increased AM symbiosis in apple roots, particularly arbuscule formation (Figure 8C and 8G). Overexpression of *MdSTR2* did not result in a significant difference in the total FA or TAG content in *MdSTR2* overexpressing roots (Figure 8H and 8I). The results indicated that MdSTR2 could affect AM symbiosis, likely by altering FA transport but not synthesis. These results supported the idea that the transcription factor MdABF2 affects the formation of AM symbiosis by regulating the expression of genes responsible for FA synthesis and translocation to the AM hyphae in apple roots.

**Figure 7.**
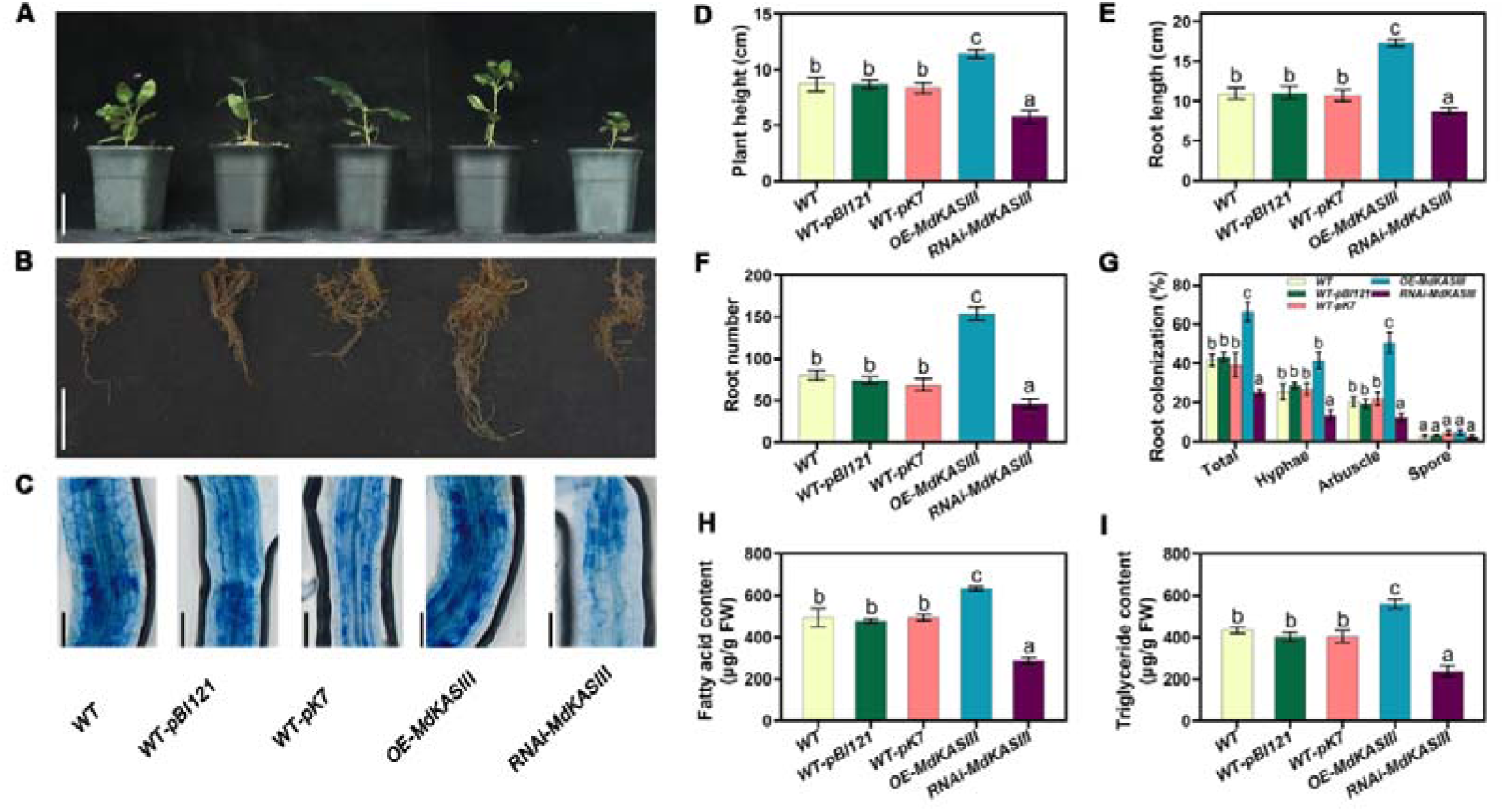
*MdKASIII* positively regulates AM symbiosis in apple (*Malus hupehensis* Rhed) seedlings. Above-ground growth phenotypes **(A)**, root growth phenotypes **(B)**, and trypan blue staining of the fungus to reveal the arbuscule morphology **(C)** of 60-day-old WT, WT-pBI121, WT-pK7, OE-*MdKASIII*, and RNAi-*MdKASIII* transgenic hairy root apple lines. Average plant height **(D)**, root length **(E)**, and root number **(F)** of transgenic hairy root apple lines. **(G)** Quantification of mycorrhizal colonization levels. **(H)** Average fatty acid content **(H)** and triglyceride content **(I)** of transgenic apple roots. **(A)** Scale bars, 5 cm. **(B)** Scale bars, 5 cm. **(C)** Scale bars, 100 µm. FW,,fresh weight. The genetic modification of the apple seedlings was only applied to the root. (**D, E**, **F**, and **G)** The bars represent the mean value ± SD (n = 9 independent biological replicates).(H and I) The bars represent the mean value ± SD (n = 3 independent biological replicates). Mixed samples from three apple seedlings carrying transgenic hairy roots were as one replicate. (D, E, F, G, H, and I) Different letters indicate significant difference (analysis of variance [ANOVA]), Duncan’s multiple range test; *P* < 0.05).

**Figure 8.**
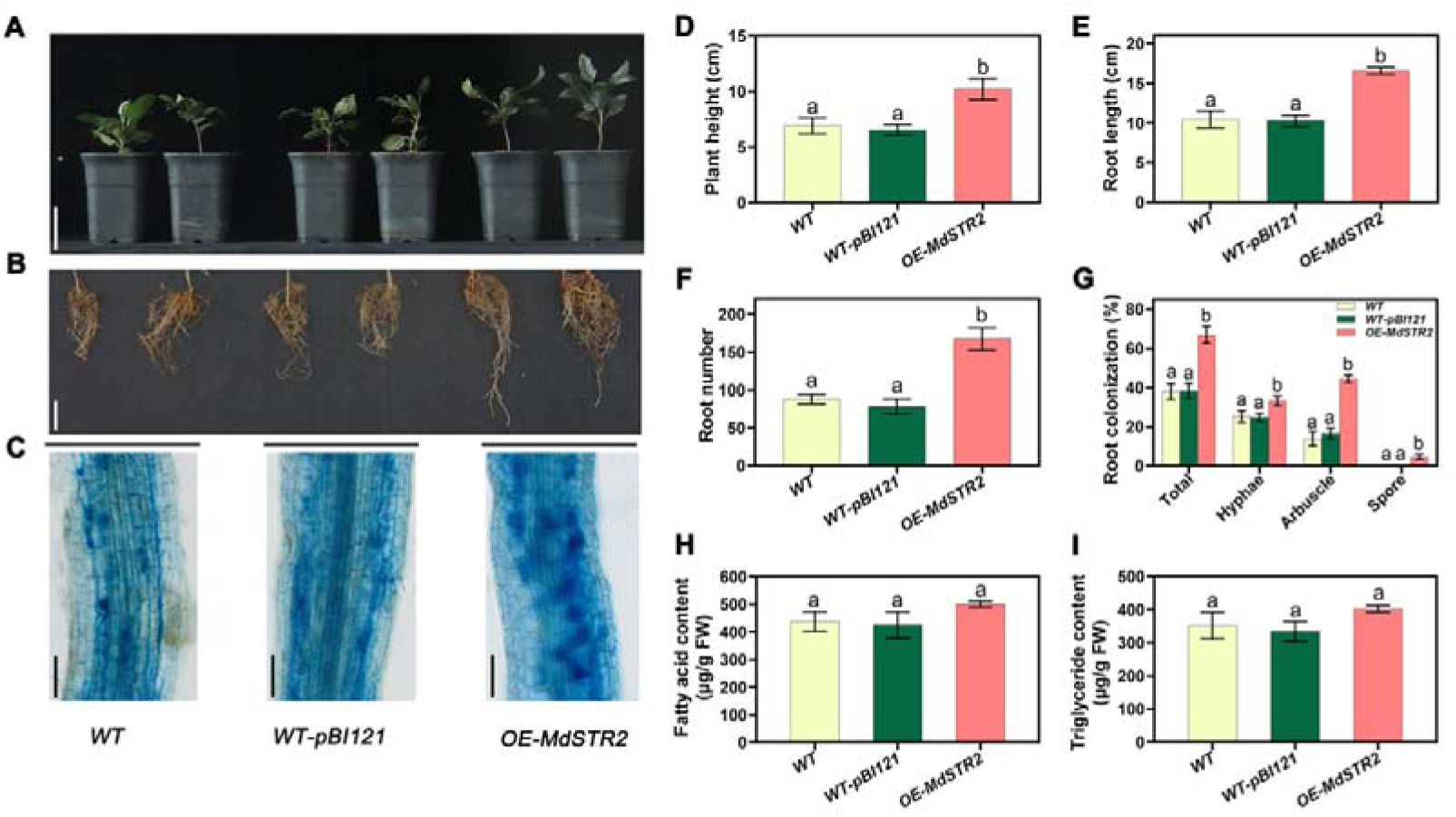
*MdSTR2* positively regulates AM symbiosis in apple (*Malus hupehensis* Rhed) seedlings. Above-ground growth phenotypes **(A)**, root growth phenotypes **(B)**, and trypan blue staining of the fungus to reveal the arbuscule morphology **(C)** of 60-day-old WT, WT-pBI121, and OE- *MdSTR2* transgenic hairy root apple lines. Average plant height **(D)**, root length **(E)**, and root number **(F)** of transgenic hairy root apple lines. **(G)** Quantification of mycorrhizal colonization levels. **(H)** Average fatty acid content **(H)** and triglyceride content **(I)** of transgenic apple roots. **(A)** Scale bars, 5 cm. **(B)** Scale bars, 5 cm. **(C)** Scale bars, 100 µm. FW, fresh weight. The genetic modification of the apple seedlings was only applied to the root. (**D, E**, **F**, and **G)** The bars represent the mean value ± SD (n = 9 independent biological replicates). **(H** and **I)** The bars represent the mean value ± SD (n = 3 independent biological replicates). Mixed samples from three apple seedlings carrying transgenic hairy roots were as one replicate. (**D, E**, **F**, **G, H,** and **I)** Different letters indicate significant difference (analysis of variance [ANOVA]), Duncan’s multiple range test; *P* < 0.05).

## Discussion

AM fungi are beneficial, symbiotic fungi that enhance nutrient uptake and stress resistance of its host plants (Zhou et al., 2015; Bernaola et al., 2018). However, the natural colonization rate of AM fungi is low in production situations (Vallino et al., 2009; Lumini et al., 2011). Therefore, it is important to enhance the symbiosis rate of AM fungi in plant roots. While low-phosphorus and nitrogen treatments have been shown to increase the infection rate, these methods are difficult to achieve in modern agricultural production (Breuillin et al., 2010; Balzergue et al., 2011; Kobae et al., 2016). Therefore, exploring the infection mechanism of AM fungi and increasing the infection rate by altering the plant itself is of great significance.

### Increased ABA synthesis is necessary for formation of AM symbiosis

ABA indeed plays an indispensable role in the development of symbiosis between apple and the fungus *Rhizophagus irregularis*. The increase in ABA during AM fungal infection is caused by endogenous ABA synthesis in the apple plant (Figure 1D; Figure 3; Supplemental Figure 3). In other words, the AM fungus either directly or indirectly triggers the plant’s ABA synthesis pathway through activating genes such as *NCED*, a key gene of ABA synthesis. It remains unknown what causes the increased expression level of *NCED*, although there are three possible pathways. First, AM fungus releases a large number of effector molecules that reprogram plant cells (Betz et al., 2024). The influence of effectors on NCED is not without precedent, and Li et al. (2020) found that effector proteins can interfere with the localization of HbNCED5 and inhibit ABA synthesis. Second, ABA synthesis is influenced by other hormones (Cutler et al., 2010; Nakashima and Yamaguchi−Shinozaki, 2013). During AM symbiosis, the levels of strigolactones (SL), salicylic acid (SA), jasmonic acid (JA), and ethylene (Eth) all change, but less so than ABA (Fig. 1C). However, as endogenous plant hormones, slight changes may cause alterations in NCED expression. Studies have shown that Eth, SA, and JA can directly or indirectly affect the expression of gene encoding key enzymes in ABA synthesis (such as NCED) (Peleg and Blumwald, 2011; Zhang et al., 2016). Third, the increase in NCED may be due to the plant’s efforts to remove excess reactive oxygen species (ROS). AM fungal invasion generates a large amount of ROS in plants (Salzer et al., 1999; Espinosa et al., 2014). To reduce oxidative damage, the plant may upregulate *NCED* expression to increase ABA levels and activate ABA signaling. In summary, the increase in NCED expression during AM fungi symbiosis may be an important regulatory mechanism for plants to cope with stress and to initiate and maintain symbiotic relationships.

Interestingly, the ten-fold increase in ABA during AM symbiosis did not inhibit plant growth or development (Figure 1). This is likely related to the suppression of the ABA signaling pathway (Santner et al., 2009). Although the ABA content significantly increased during the AM fungal infection process, the downstream ABA signal transduction genes *PYL* and *SnRK* did not show a significant difference before and after inoculation (Supplemental Data 1). Additionally, there is often a homeostatic balance among hormones during plant growth and development (Depuydt and Hardtke, 2011; Vanstraelen and Hardtke, 2012). Although the content of ABA significantly increases, the levels of the growth-promoting hormones GA, SL, and BL also significantly rise during AM fungal infection (Figure 1C).

### ABA regulated the synthesis and transport of FA to provide carbon for AM fungal cells in roots

During AM symbiosis, fatty acids are a crucial nutrient source for AM fungi. Genes related to FA synthesis, such as *KAS*, *FatM*, and *RAM*, are induced by AM fungi, and overexpression of these genes can also promote mycorrhizal symbiosis (Jiang et al., 2017). In apple, ABA affected AM symbiosis by influencing the synthesis and transport of FA (Figure 5, Figure 6, Figure 7 and Figure 8).

The regulatory hierarchy centered around ABF that controls the direct FA metabolism pathway appears to be conserved across different plant species (Yeap et al., 2017; Li et al., 2020). Numerous studies have also shown that ABA is closely related to FA: in oil palm, EgABI5 transcriptionally activates *EgDGAT1*, promoting oil biosynthesis during fruit development (Yeap et al., 2017). In our study, ABA, through the transcription factor ABF2, directly or indirectly regulates genes related to synthesis (*KASI* and *KASIII*) and transport (*STR2*) of FA, thereby affecting the supply of FA and influencing AM symbiosis (Figure 4, Figure 5, Figure 6, Figure 7 and Figure 8). This indicates that ABF2 is crucial for AM symbiosis.

In *Medicago sativa*, the AP2-domain transcription factor WRI5a is the main regulatory factor for lipid biosynthesis and transfer during AM symbiosis, and WRI5a could regulate STR expression (Jiang et al., 2018). We found that ABF2 could activate the expression of *MdWRI3* (a homolog of *WRI5a*). ABF2 can directly regulate the expression of *STR2* or indirectly regulate *STR2* expression through *WRI3*. STR controls the lipid flux from the plant to the AM fungus (Zhang et al., 2010; Gutjahr et al., 2012; Jiang et al., 2017).

It is interesting that plants have two regulating pathways to transfer FA to AM fungi. In fact, we found that multiple genes related to FA synthesis, such as *RAM2* and *RAM2-1*, and *KAS1*, *KAS1-1*, *KASIII*, and *KASIII*, were significantly upregulated (Supplemental Figure 9), leading to a significant increase in FA content, especially TAG (Figure 7). Comparatively, only one gene related to FA transfer, STR2, was upregulated (Supplemental Figure 9). Given our findings that ABF2 both directly and indirectly regulates STR2 expression (Figure 6), we propose that AM fungi may have a higher demand for FA compared to other carbon compounds in apple roots; the sharp increase in FA content in the mycorrhiza requires the plant to open the transfer pathways to meet AM fungus’s needs.

Overall, our research presents significant progress in understanding the molecular mechanisms by which ABA promotes AM fungal colonization during the complex interaction between plants and fungi. The results showed that ABA is necessary to efficiently form AM symbiosis in apple root. The infection of AM fungus could induce ABA synthesis, with upregulated expression of *MdNCED3.1/3.2* in the root via an unknown pathway. This increased ABA content upregulated the expression of genes related to both FA synthesis (e.g., *MdKASIII*) and FA transport (*MdSTR2*) through the MdABF2 signaling pathway. Increased levels of FAs from the plant roots are a primary carbon supply for fungal growth, thereby promoting AM symbiosis (Figure 9). Other studies have also shown that ABA synthesis significantly increases during AM symbiosis. One way to improve AM symbiosis of crops, with the aim of enhancing crop yield and fruit quality, may be increasing ABA levels in roots. Further exploration of the specific mechanisms by which AM symbiosis affects plant metabolism and development is both highly necessary and highly feasible in our pursuit of better utilization of AM fungi to improve crop yield and quality.

**Figure 9.**
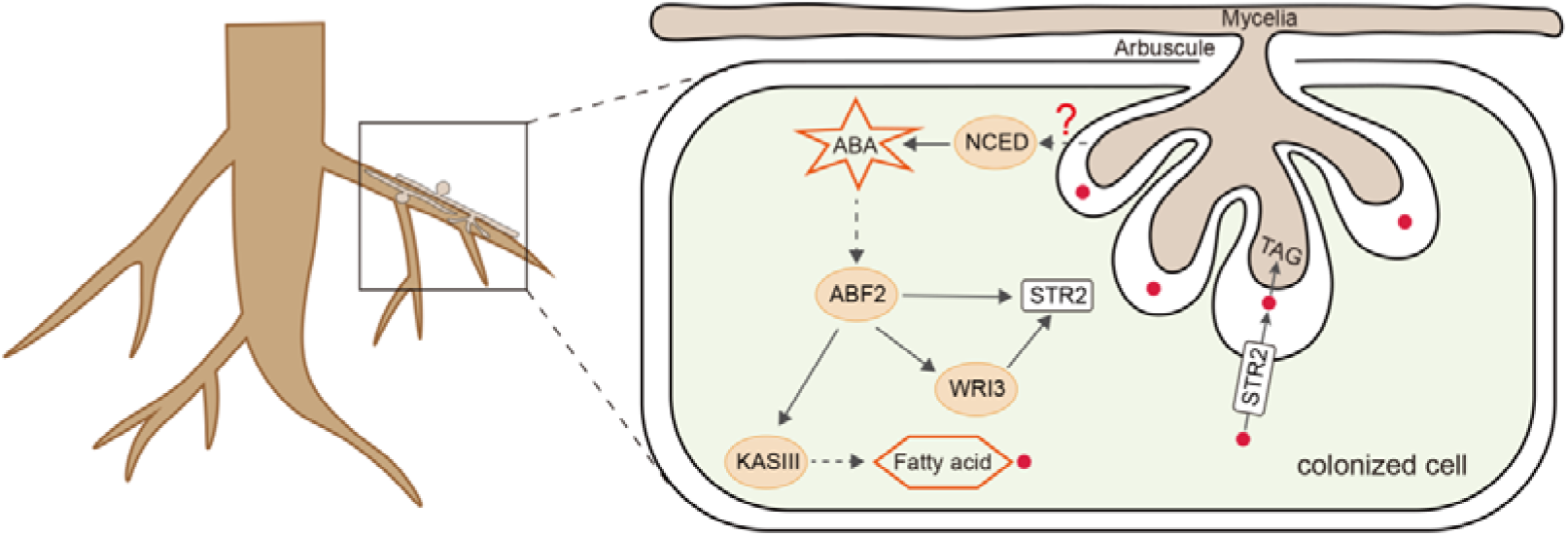
Proposed pathway through which ABA regulates the supplementation of fatty acids to arbuscular mycorrhizal fungus during symbiosis in root cells. The infection of AM fungus induces ABA synthesis through upregulated expression of *MdNCED3.1/3.2* in the root via an unknown pathway. The higher ABA level then increases the expression of genes that synthesize (e.g., *MdKASIII*) and transport (*MdSTR2*) fatty acids through the MdABF2-mediated transduction pathway. Unknown lipid transport proteins in the AM fungus further absorb the lipids, resulting in a carbon supply from the plant roots necessary for fungal growth, thereby promoting AM symbiosis. TAG, triacylglycerol.

## Materials and methods

### Plant and fungal materials

This experiment was carried out in Shaanxi Province at the Northwest A&F University. The plants were maintained in the greenhouse in plastic pots filled with sand (sterilized by steaming at 121°C for 2 h). The AM fungus Rhizophagus irregularis (line number: BGC BJ09) was maintained in a 4°C refrigerator. Spores were isolated and counted as described (Gerdemann and Nicolson, 1963). The inoculant contained about 60 spores per gram.

Apple rootstock cultivar M26 (*Malus pumila* Mill.) plants and *Malus hupehensis* Rhed seedlings were used in these experiments. M26 was used to study the effect of exogenous ABA treatment on AM symbiosis and was also used for omics sequencing. *Malus hupehensis* Rhed, as a triploid,is typical apomixis-type of *Malus* species, and was used for its susceptibility to hairy root transformation by *Agrobacterium rhizogenes*.

### Experimental treatments

For all experiments, the roots of apple plants/seedlings were inoculated with 20 g of AM fungal inoculant, while the control group was inoculated with 20 g of sterilized AM fungal spores. The plants were placed in the greenhouse after treatment at a relative humidity between 55 – 75% and temperatures between 12–30 °C). The plants were watered normally, and all plants were watered weekly with 100 mL ½-strength low-phosphorus Hoagland’s nutrient solution.

For transcriptome sequencing and metabolic analyses, 200 saplings of apple rootstock M26 were used, with 100 plants in the treatment group and 100 in the control group. The plants were about 10 cm in height and were grown in plastic pots (21 cm × 15 cm). Sixty days after inoculation with AM fungi, the plants were measured and root samples were collected.

The apple rootstock M26 was used for exogenous ABA treatment and planted in plastic pots (14 cm × 12 cm). Plants about 10 cm in height were selected. A quarter of them were left un-inoculation and untreated for 45 days and considered a control group. A quarter of them were inoculated with AM fungal spores and allowed to grow for 45 days. Another quarter of them were subjected to fungal inoculation, but after just 30 days they were treated with exogenous ABA (50 µmol/L) and allowed to grow for another 15 days. A fourth experimental group of plants were given exogenous Flu (50 µmol/L) 30 days after fungal inoculation before sampling 15 days later. The plant indexes were measured, and samples were collected 45 days after AM fungal inoculation.

The apple seedlings (*Malus hupehensis* Rhed) were used for generation of *MdNCED3.1/3.2* transgenic hairy roots and were planted in plastic pots (8.8 cm × 9 cm). One part was inoculated with AM fungal spores, the other part was not inoculated. Plants in these groups were not transformed (WT). And others were co-cultured with *Agrobacterium rhizogenes* harboring either pBI121 empty vector, pK7 empty vector, *pBI121-MdNCED3.1*, *pBI121-MdNCED3.2*, *pK7-MdNCED3.1*, or *pK7-MdNCED3.2*. The transgenic lines that had not been inoculated were split into 7 groups as the control group. Another set of inoculated lines were used as experimental groups as above. After hairy root transformation with *Agrobacterium rhizogenes*, the plants were cultured for 60 days before phenotypic determination and sample collection.

The *MdABF2*, *MdKASIII*, and *MdSTR2* transgenic materials were developed and treated in the same way as the *MdNCED3.1*/*3.2* roots.

### Mycorrhizal colonization and morphological index assays

The method used to observe mycorrhizal colonization was described by He et al. (2016), with minor modifications. The roots were cleaned, soaked in 10% KOH until transparent and softened, stained with 0.05% trypan blue, and decolorized 3 times with 1:1 acetic acid-glycerin before microscopic examination. At the end of the 60-day co-culture period, plant height and root length were measured using a steel ruler, and the number of roots were manually counted.

### Transcriptome analysis of apple roots

The root tips were used to extract RNA according to the Plant Total RNA Isolation Kit (FOREGENE) protocol. Primers sequences are shown in Supplemental Table 1. A high-quality cDNA library was constructed. The transcriptome sequence was performed on an Illumina HiSeq platform (Illumina, San Diego, CA, USA). Raw data in fast format (original reads) was processed by an internal Perl script. After program processing, high-quality reads were obtained. High-quality reads were mapped to the reference genome (https://www.rosaceae.org/species/malus/all), and the FPKM value of each gene was calculated. We utilized PCA to evaluate the connections among the samples. Differentially expressed genes (DEGs) were screened based on a P-value <0.05 and a fold change ≥2.0. The KEGG orthogonal and MapMan databases were used to annotate functions of the DEGs.

### Hormone analysis in apple roots

Hormone metabolomics were used to identify differential hormone levels by UPLC-MS/MS analysis. Briefly, 0.05 g of root tip samples were mixed with 10 µl of a mixed internal standard solution (10 ng/ml), and 1 ml of methanol, water, and formic acid (15:4:1, v/v/v) and mixed well by vortex. A Thermo Vanquish UPLC system with Waters ACQUITY UPLC HSS T3 C18 column (1.8 µm, 100 mm×2.1 mm i.d.) was used. The program was set as follows: a flow rate of 0.35 mL/min, a column temperature of 40 °C, and an injection volume of 4 µL each time. Solvents A and B were water/0.04% acetic acid and ACN/0.04% acetic acid, respectively. For mass spectrometry (MS) analysis, the electrospray ionization (ESI) temperature was 550 °C and the curtain gas (CUR) pressure was 35 psi. PCA was used to assess the relationships between samples. Differential hormone metabolite levels were screened based on a P-value <0.05 and a fold change ≥2.0.

### Hairy Root Agrobacterium Transformation, Mycorrhizal Infestation

To create overexpression and RNA interference vectors, the coding sequences of MdNCED3.1, MdNCED3.2, MdABF2, MdKASIII, and MdSTR2 were inserted into a pBI121 vector containing the 35S promoter. Additionally, specific truncated sequences were added to a pK7GWIWG2 vector. Next, *Agrobacterium rhizogenes* K599 (Weidi Biotechnology, Shanghai, China) was transformed using the fusion vectors, pBI121-GFP and pK7GWIWG2.

We created transgenic plants using *Agrobacterium rhizogenes* to generate hairy roots in a way similar to what Boisson-Dernier (2001) et al. described. Sixty days after co-culture with the *A. rhizogenes*, the roots were co-cultured with *Rhizophagus irregularis* spores according to Jiang et al. (2017).

To transform apple roots, briefly, apple seeds were placed in petri dishes lined with wet filter paper and allowed to germinate. After the seedlings had three cotyledons, the *Agrobacterium rhizogenes* strain Arqua-1 containing the desired vector was used to transform and generate hairy roots. Each vector contained a GFP (Green Fluorescent Protein) reporter gene.

After 3 to 4 weeks, untransformed roots were removed based on fluorescence of the roots in the somatic fluorescence microscope (ZEISS). The apple seedlings with transformed roots and the appropriate control plants (untransformed wild type or plants harboring empty vectors) were transferred to sand, inoculated with about 60 irregular mycospores per plant, and cultivated in the greenhouse.

### Quantification of FAs

The triglyceride content in the roots was extracted according to Li et al. (2017). And the fatty acid content in the roots was extracted according to Shi et al. (2018). Fatty acid and triglyceride analysis was performed using an Agilent 7890B gas chromatograph that had an HP-88 column (RESTEK; 0.25 mm inner diameter, 100 m length, and 0.250μm film thickness; Agilent, Shanghai, China).

Fatty acid was quantified by use of the external standard method. Using the fatty acid methyl ester mixed standard (GLC NESTLE 37 MIX, Solarbio), the fatty acid methyl ester standard curve was created.

### Quantification of ABA

ABA was extracted using the ethyl acetate method, and its content was determined using high-performance liquid chromatography (Müller and Munné-Bosch, 2011). A high-performance liquid chromatography-mass spectrometry (HPLC-MS; QTRAP5500, AB SCIEX, USA) was used with a packed column of 150 mm × 4.6 mm, 5 μm (InsertContinustTM AQ-C18, Shimadzu, Japan). Mobile phase A was 0.1% formic acid (85178, Thermo, USA), and mobile phase B was methanol (67561, Thermo, USA).

### Gene expression analysis

Roots were used for RNA extraction using the Plant Total RNA Isolation Kit (FOREGENE). Supplemental Data 3 lists all of the primers used. For qRT-PCR reactions, cDNA was synthesized with PrimeScript RT reagent (Takara, http://www.takarabiomed.com.cn). The 2x Fast qPCR Master Mixture (DiNing, https://di-ning.com.cn/) was used for cDNA analysis on the iQ5 multicolor real-time Polymerase Chain Reaction Detection System (Bio-Rad). Data were analyzed using iQ5 2.0 standard optical system analysis software and the 2^−ΔΔ^CT method (Livak and Schmittgen 2001).

### Phylogenetic analysis

To discover the potential NCED genes in the apple genome, the known NCED genes of Arabidopsis were used as a query for TBLASTN of the apple genome. We constructed multiple comparisons of the amino acid sequences of the MdNCEDs using MEGA7 (Bootstrap = 1000).

### Yeast One-hybrid (Y1H) assay

The Matchmaker Gold Y1H system was used for yeast (*S. cerevisiae*) one-hybrid assays (Takara, https://www.takarabiomed.com.cn). The reporter strains were created by putting the *MdKASI*, *MdKASIII*, *MdRAM2*, and *MdSTR2* promoters into pAbAi vectors for transformation into YIH Gold. On SD/-Ura plates with different amounts of aureomycin A (AbA), the lowest concentration to inhibit growth was found for each strain. The *MdABF2* full-length coding sequence was inserted into the pGADT7-Rec vector. The MdABF2-pGADT7-Rec was transformed into each promoter-reporter strain according to the method described in the manufacturer’s protocol. The transformed yeast cells were spread on SD/-Leu/ABA plates and incubated at 30 °C for 3 days.

### GUS staining

The promoter for *MdKASIII*, *MdWRI3*, and *MdSTR2* were inserted into the pC0390-GUS vector, respectively, while the ORF sequences sequence of MdABF2, and MdWRI3 were inserted into the pBI121, respectively (overexpression vector). The fusion vectors were transformed into *Agrobacterium rhizogenes* GV3101 (pSoup-p19) (Weidi Biotechnology, Shanghai, China).

The 35S:MdABF2 was simultaneously injected into tobacco leaves with either the *proMdKASIII-GUS*, *proMdSTR2-GUS*, or *proMdWRI3-GUS*. Likewise, the 35S:MdWRI3 was simultaneously injected into tobacco leaves with the *proMdSTR2-GUS*.Histochemical staining was performed to detect GUS activity in transformants (Zhu et al., 2023). Three biological replicates were performed for each treatment. These primers are listed in Supplemental Data 3.

### Luciferase complementation assay

The ORF sequences of *MdABF2* and *MdWRI3* were cloned into the pGreenII 62-SK vector, respectively. The promoters of *MdKASIII*, *MdKSTR2*, and *MdWRI3* were cloned into the pGreenII 0800-LUC vector, respectively. The fusion vectors were transformed into *Agrobacterium rhizogenes* GV3101 (pSoup-p19) (Weidi Biotechnology, Shanghai, China). The *A. rhi*zogenes strains were co-infiltrated into tobacco leaves in different pairings. The bioluminescence intensity was detected using a live plant imaging system (PlantView100, Guangzhou Biolight Biotechnology Co., China).

### EMSA assay

Purified MdABF2-His and MdWRI3-His recombinant proteins were produced following the directions for Ni-NTA binding resin (7 Sea Biotech, Shanghai, China). Promoter probes were amplified using primers with biotin-labeled oligonucleotides (Invitgen, Shanghai, China) or unlabeled oligonucleotides (an unlabeled competitor), as shown in Supplemental Data 3. EMSA reactions were performed as described by Zheng et al. (2018). Each experiment was independently repeated three times.

### Statistical analysis of data

Data was analyzed with GraphPad Prism 10.1.2 and IBM SPSS Statistics 26. Significant differences were determined using one-way analysis of variance (ANOVA) with two-sided Student’s *t*-test or Duncan’s multiple range test (*p* < 0.05).

## Acknowledgments

c

## Author contributions

M.J.L, M.R.Z, and F.W.M.: conceived and supervised this study; S.J., C.H.L. and L.J.D.: performed the experiments; Y.C.L., L.C.Z., J. S. and X.Y.W.: performed the bioinformatics analysis; S.J.: wrote the manuscript; M.J.L, B.Q.M, B.Q.M, Y-L.R, M.R.Z and S.J.: discussed the study and revised the manuscript.

The authors responsible for distribution of materials integral to the findings presented in this article in accordance with the policy described in the Instructions for Authors are: ManRang Zhang, Mingjun Li and Fengwang Ma

---

**Supplemental Figure 1.**
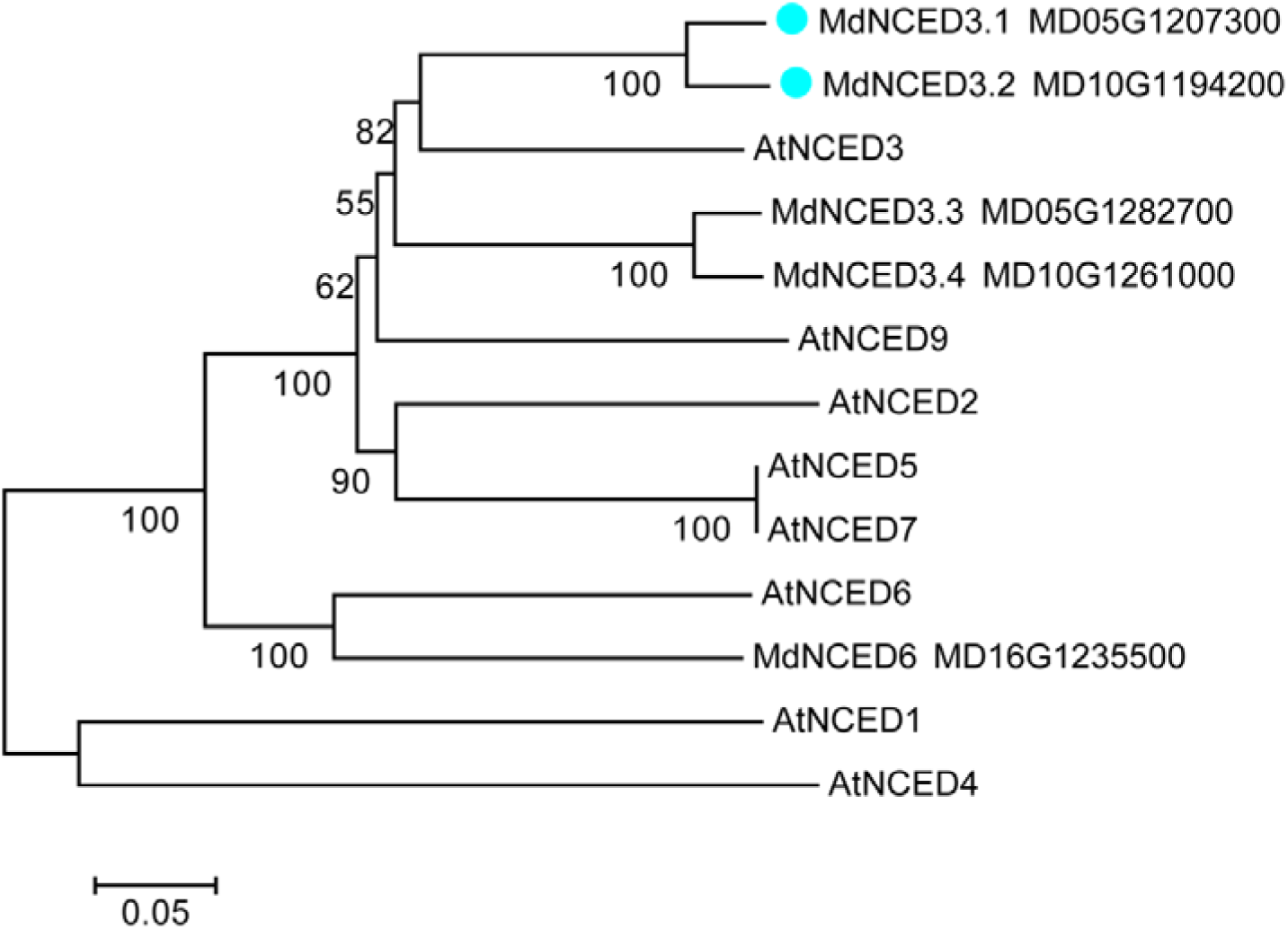
Phylogenetic analysis of 9-cis-epoxycarotenoid dioxygenase (NCED) proteins from apple (*Malus* × *domestica*) and *Arabidopsis thaliana*. The phylogenetic tree was constructed using the maximum likelihood method of the MEGA7 software. Bootstrap analysis of 1000 trials provided a reliable estimate of the topology of the phylogenetic tree.

**Supplemental Figure 2.**
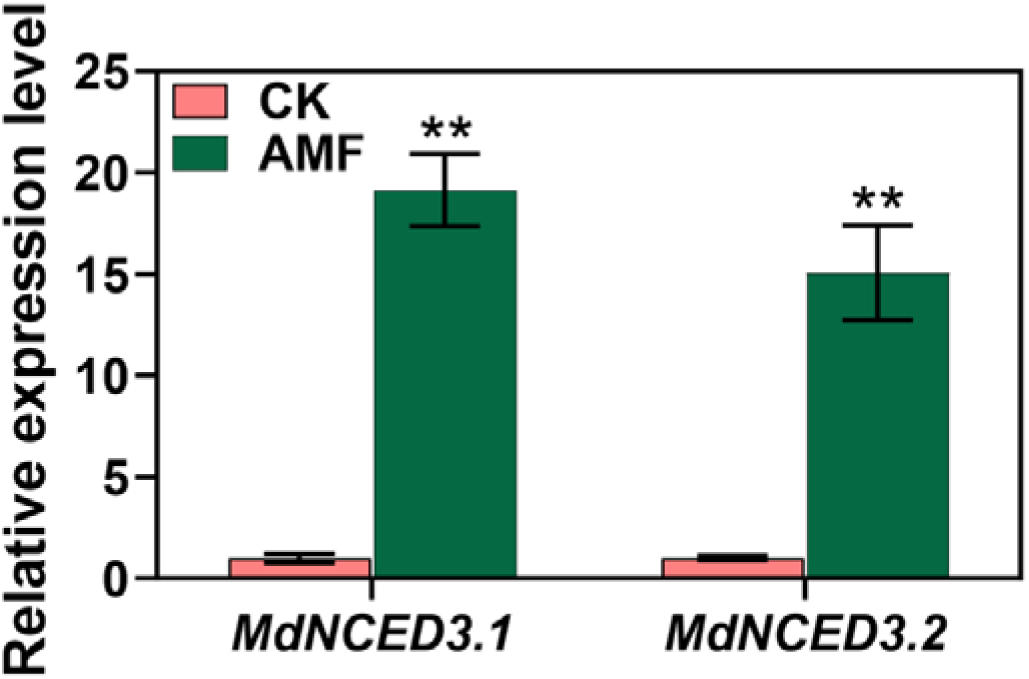
The relative expression levels of *MdNCED3.1* and *MdNCED3.2* in the roots of 60-day-old uninoculated M26 (*Malus pumila* Mill.) plants and M26 plants inoculated with AMF. The transcript levels were normalized to those of *MdActin*. Relative expression levels for each gene were obtained via the ddCT method, with its expression in uninoculated M26 plants set as ‘1’. The bars represent the mean value ± SD (n = 3 independent biological replicates). Mixed samples from three M26 plants were as one replicate. The asterisks indicate significant differences as assessed by one-way ANOVA (two-sided Student’s *t*-test; \*\**P* < 0.01).

**Supplemental Figure 3.**
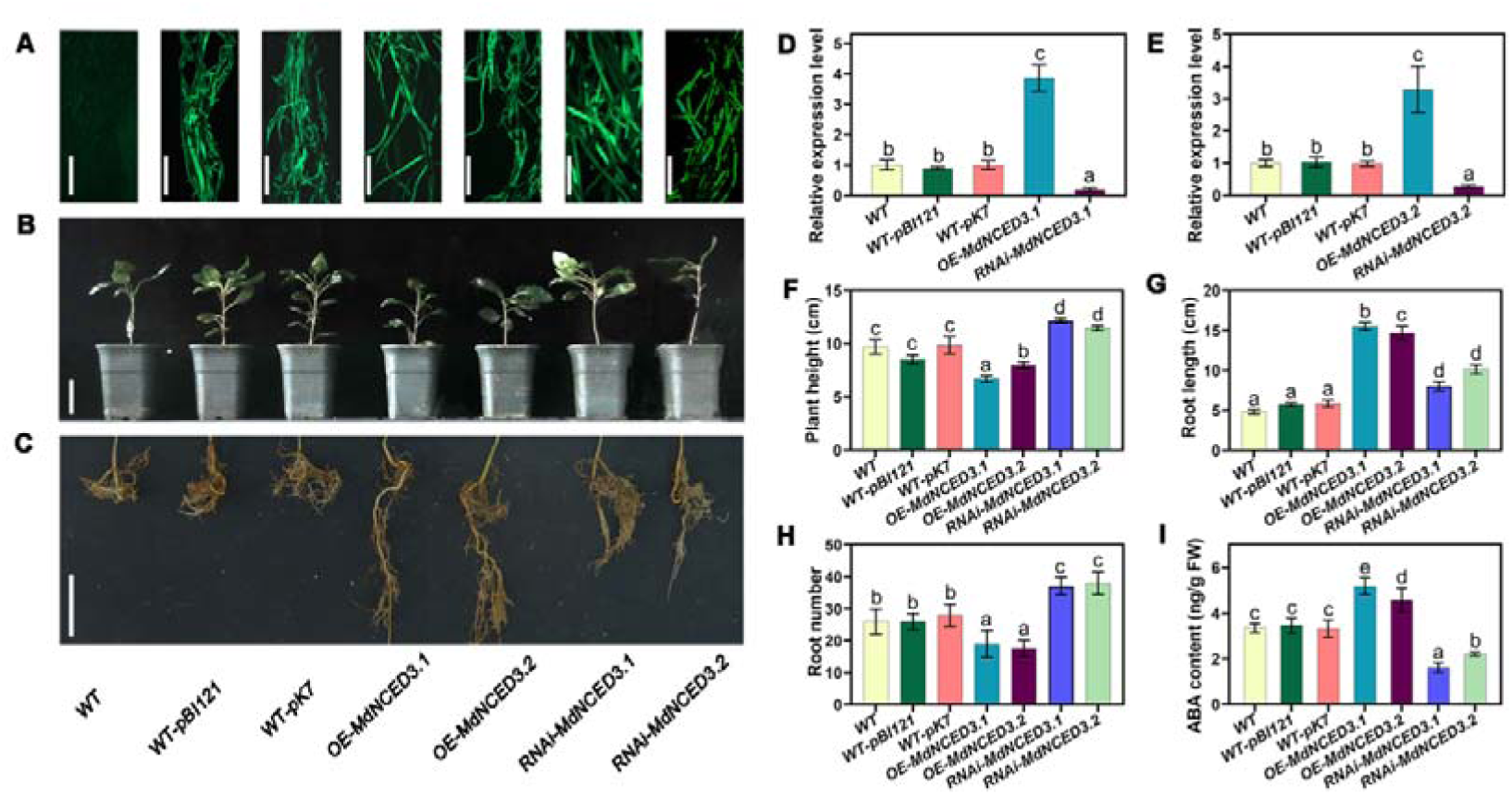
The effects of changing the expression of *MdNCED3.1*/*3.2* in the roots of 60-day-old apple (*Malus hupehensis* Rhed) seedlings without inoculated AMF. Each construct contained a GFP cassette for screening of transformation. **(A)** Images of the transgenic root systems of apple seedlings with green fluorescent protein. Scale bars, 1 mm. **(B)** Above-ground growth phenotypes of transgenic hairy root apple lines. Scale bars, 5 cm. **(C)** Root growth phenotypes of transgenic hairy root apple lines. Scale bars, 5 cm. **(D)** Relative expression levels of *MdNCED3.1* mRNA in transgenic apple roots to the WT control (set as ‘1’). **(E)** Relative expression levels of *MdNCED3.2* mRNA in transgenic apple roots compared to the WT control (set as ‘1’). **(F)** Plant height of apple seedlings carrying transgenic hairy roots. **(G)** Root length of apple transgenic lines. **(H)** Root number of untransformed and transgenic apple lines carrying a hairy root construct. **(I)** The ABA content in the root of apple transgenic lines. FW, fresh weight; WT, wild type; WT-pBI121, Apple seedlings transformed with an hairy root empty vector for overexpression and containing the GFP tag (plasmid Binary Vector 121); WT-pK7, Apple seedlings transformed with an RNA interference empty vector containing the GFP tag (pK7GWIWG2); OE-*MdNCED3.1*, *MdNCED3.1*-overexpressing root lines; OE-*MdNCED3.2*, *MdNCED3.2*-overexpressing root lines; RNAi-*MdNCED3.1*, *MdNCED3.1*-RNA interference root lines; RNAi-*MdNCED3.2*, *MdNCED3.2*-RNA interference root lines. **(D**, **E**, and **I)** The bars represent the mean value ± SD (n = 3 independent biological replicates). Samples from three carrying transgenic hairy roots were as one replicate. (**F**, **G,** and **H)** The bars represent the mean value ± SD (n = 9 independent biological replicates). (**D, E**, **F**, **G, H,** and **I)** Different letters indicate significant difference (analysis of variance [ANOVA]), Duncan’s multiple range test; *P* < 0.05).

**Supplemental Figure 4.**
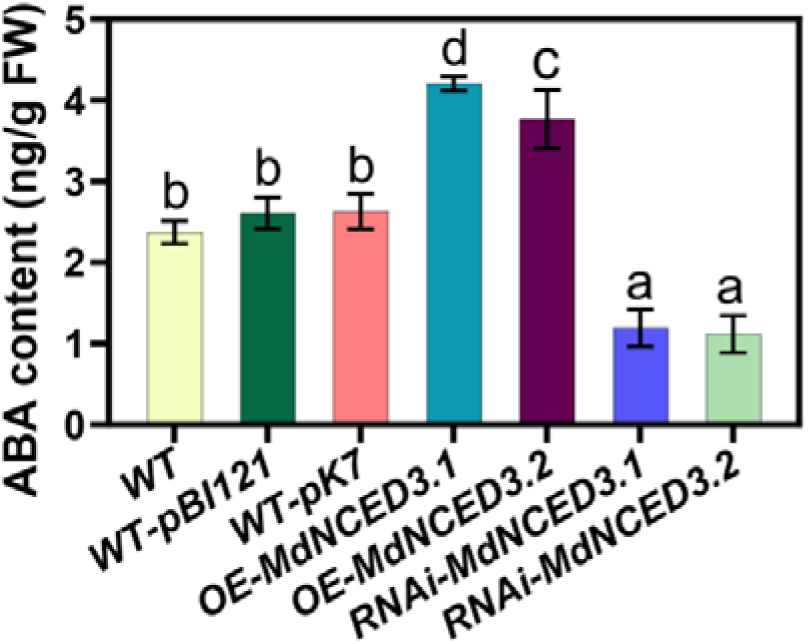
The ABA content with *MdNCED3.1/3.2* transgenic hairy roots in apple root. FW, fresh weight; WT, wild type; WT-pBI121, Apple seedlings transformed with an overexpressed empty vector containing the GFP tag (plasmid Binary Vector 121); WT-pK7, Apple seedlings transformed with RNA interference empty vector containing the GFP tag (pK7GWIWG2); *OE-MdNCED3.1*, MdNCED3.1-overexpressing root lines; *OE-MdNCED3.2*, MdNCED3.2-overexpressing root lines; *RNAi-MdNCED3.1*, MdNCED3.1-RNA interference root lines; *RNAi-MdNCED3.2*, MdNCED3.2-RNA interference root lines. The bars represent the mean value ± SD (n = 3 independent biological replicates). Mixed samples from three apple seedlings carrying transgenic hairy roots were as one replicate. Different letters indicate significant difference (analysis of variance [ANOVA]), Duncan’s multiple range test; *P* < 0.05).

**Supplemental Figure 5.**
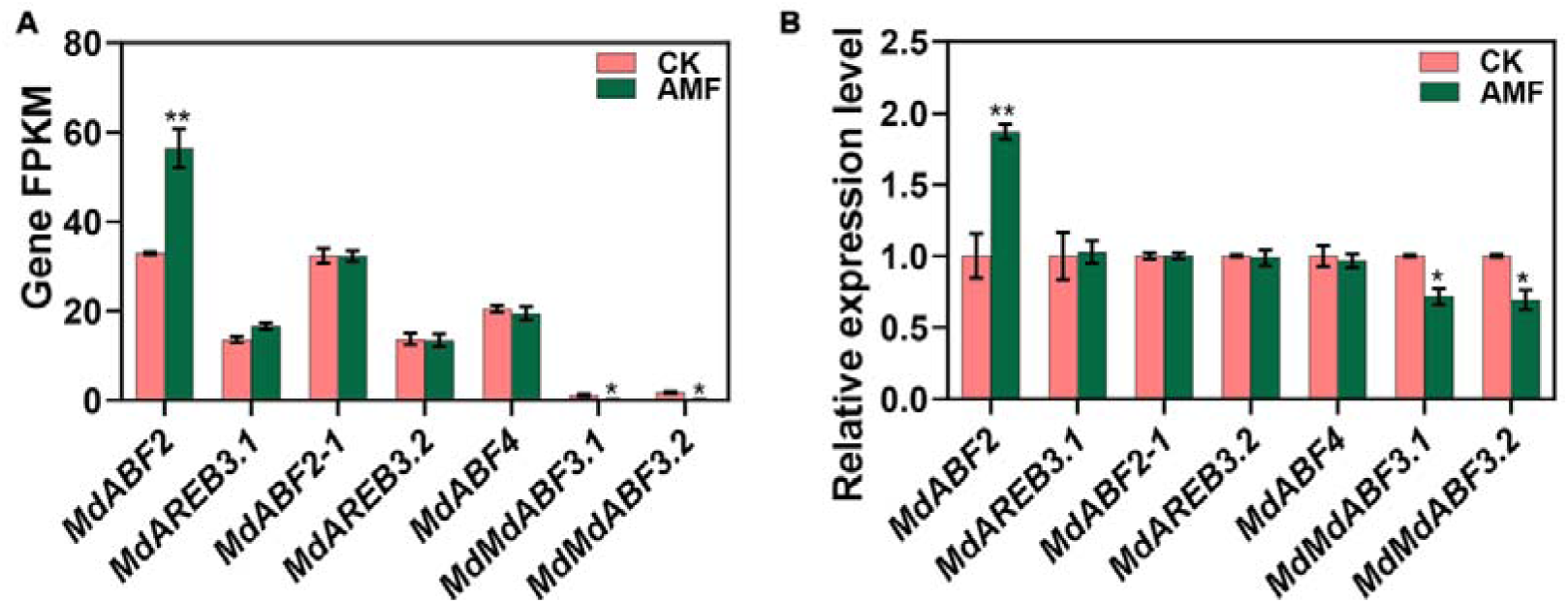
**(A)** RNAseq-based FPKM levels of *MdABF*/*MdAREB* expression in the roots of 60-day-old inoculated M26 (*Malus pumila* Mill.) plants were compared to those in uninoculated M26 plants. **(B)** The qPCR-based relative expression levels of *MdABF*/*MdAREB* in the roots of 60-day-old inoculated M26 (*Malus pumila* Mill.) plants were compared to those in uninoculated M26 plants. The transcript levels were normalized to those of *MdActin*. Relative expression levels for each gene were obtained via the ddCT method, with its expression in uninoculated M26 plants set as ‘1’. The bars represent the mean value ± SD (n = 3 independent biological replicates). Mixed samples from three M26 plants were as one replicate. The asterisks indicate significant differences as assessed by one-way ANOVA (two-sided Student’s *t*-test; \*\**P* < 0.01, \**P* < 0.05).

**Supplemental Figure 6.**
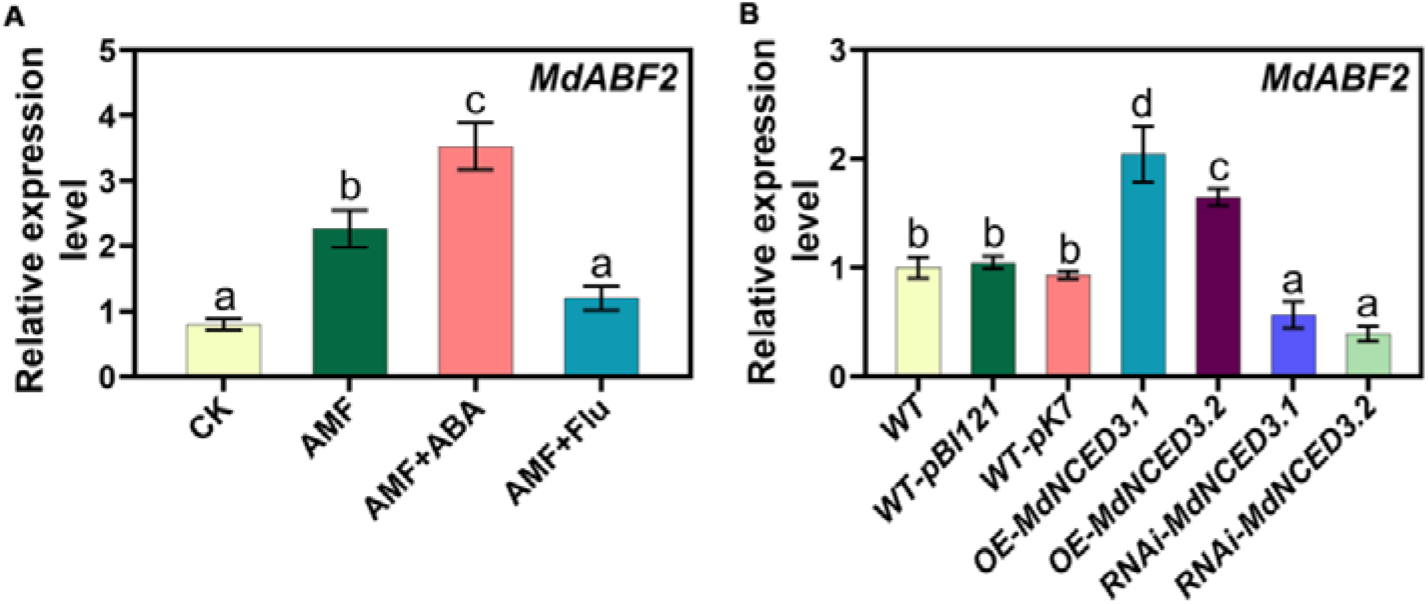
Relative expression levels of *MdABF2* in apple roots following *R*. *irregularis* infection. **(A)** Relative expression levels of *MdABF2* in the roots of 45-day-old M26 (*Malus pumila* Mill.) plants under CK, AMF, AMF+ABA, and AMF+Flu conditions, as determined by qPCR, compared to the CK (set as ‘1’). CK, Un-inoculated plants without ABA treatment for 45 days as control. AMF, Mycorrhizal-inoculated plants without ABA treatment for 45 days. AMF+ABA, Mycorrhizal-inoculated plants were grown for 30 days before spraying with exogenous ABA (50 µmol/L) and further growth for 15 days. AMF+ Flu, mycorrhizal-inoculated plants were grown for 30 days before spraying with exogenous Flu (50 µmol/L ABA synthesis inhibitor fluoridone) and further growth for 15 days. The bars represent the mean value ± SD (n = 3 independent biological replicates). Samples from 3 M26 plants were as 1 replicate. **(B)** The relative expression levels, based on qPCR, of MdABF2 (*Malus hupehensis* Rhed) in the roots of 60-day-old WT-pBI121, WT-pK7, *OE-MdNCED3.1*, *OE-MdNCED3.2*, *RNAi-MdNCED3.1*, and *RNAi-MdNCED3.2* apple seedlings with transgenic hairy roots, compared to untransformed WT controls (set as ‘1’). The bars represent the mean value ± SD (n = 3 independent biological replicates). Mixed samples from three apple seedlings carrying transgenic hairy roots were as one replicate. The transcript levels were normalized to those of *MdActin*. Relative expression levels for each gene were obtained via the ddCT method. Different letters indicate significant difference (analysis of variance [ANOVA]), Duncan’s multiple range test; *P* < 0.05).

**Supplemental Figure 7.**
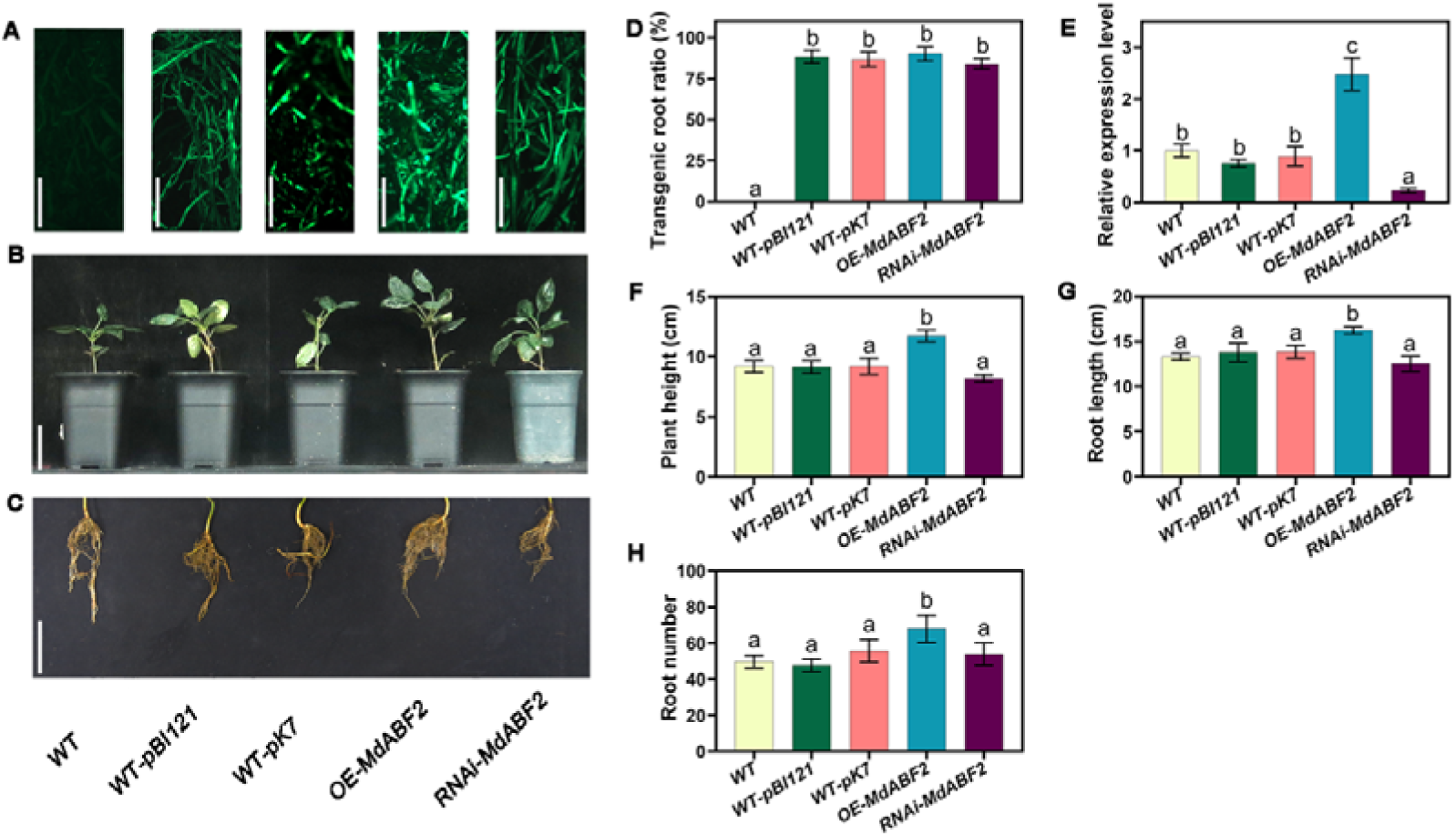
The effects of changing the expression of *MdABF2* in the roots of 60-day-old apple (*Malus hupehensis* Rhed) seedlings not inoculated with AMF. **(A)** Images of the transgenic root systems of apple seedlings expressing green fluorescent protein. Scale bars, 1 mm. **(B)** Above-ground growth phenotypes of apple seedlings carrying transgenic hairy roots. Scale bars, 5 cm. **c** Root growth phenotypes of apple seedlings carrying transgenic hairy roots. Scale bars, 5 cm. **(D)** Transgenic root ratio of apple seedlings carrying transgenic hairy roots after root co-culture with *Agrobacterium rhizogenes*. **(E)** The relative expression levels of *MdABF2* mRNA in the root of the apple seedlings carrying transgenic hairy roots with the indicated constructs compared to the WT control (set as ‘1’). The bars represent the mean value ± SD (n = 3 independent biological replicates). Mixed samples from three apple seedlings carrying transgenic hairy roots were as one replicate. **(F)** Plant height of apple seedlings carrying transgenic hairy roots. **(G)** Root length of apple seedlings carrying transgenic hairy roots. **(H)** Root number of apple seedlings carrying transgenic hairy roots. WT, wild type; WT-pBI121, apple seedlings transformed with an empty overexpression vector containing the GFP tag (plasmid Binary Vector 121); WT-pK7, apple seedlings transformed with an empty RNA interference vector containing the GFP tag (pK7GWIWG2); OE-*MdABF2*, *MdABF2*-overexpressing root lines; RNAi-*MdABF2*, *MdABF2*-RNA interference root lines. (**D**, **F**, **G**, and **H)** The bars represent the mean value ± SD (n = 9 independent biological replicates). ((**D**, **E**, **F**, **G**, and **H)** Different letters indicate significant difference (analysis of variance [ANOVA]), Duncan’s multiple range test; *P* < 0.05).

**Supplemental Figure 8.**
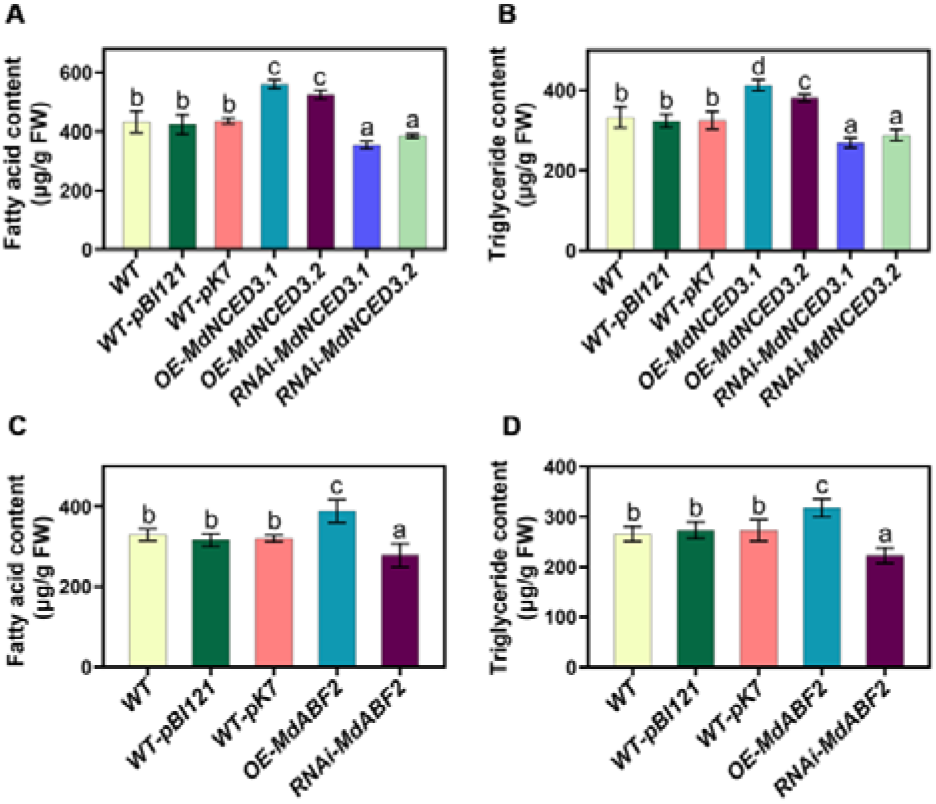
Fatty acid and triglyceride content in the roots of 60-day-old apple (*Malus hupehensis* Rhed) seedlings carrying transgenic hairy roots without *R*. *irregularis* infection of the roots. **(A), (B)** Fatty acid (**A)** and triglyceride (**B)** content of WT, WT-pBI121, WT-pK7, OE-MdNCED3.1, OE-MdNCED3.2, RNAi-MdNCED3.1, and RNAi-MdNCED3.2 apple seedlings with transgenic hairy roots, compared to untransformed WT controls. **(C), (D)** Fatty acid **(C)** and triglyceride (**D)** content of WT, WT-pBI121, WT-pK7, *OE-MdABF2*, and *RNAi-MdABF2* apple transgenic hairy roots. FW, fresh weight. The bars represent the mean value ± SD (n = 3 independent biological replicates). Mixed samples from three apple seedlings carrying transgenic hairy roots were as one replicate. Different letters indicate significant difference (analysis of variance [ANOVA]), Duncan’s multiple range test; *P* < 0.05).

**Supplemental Figure 9.**
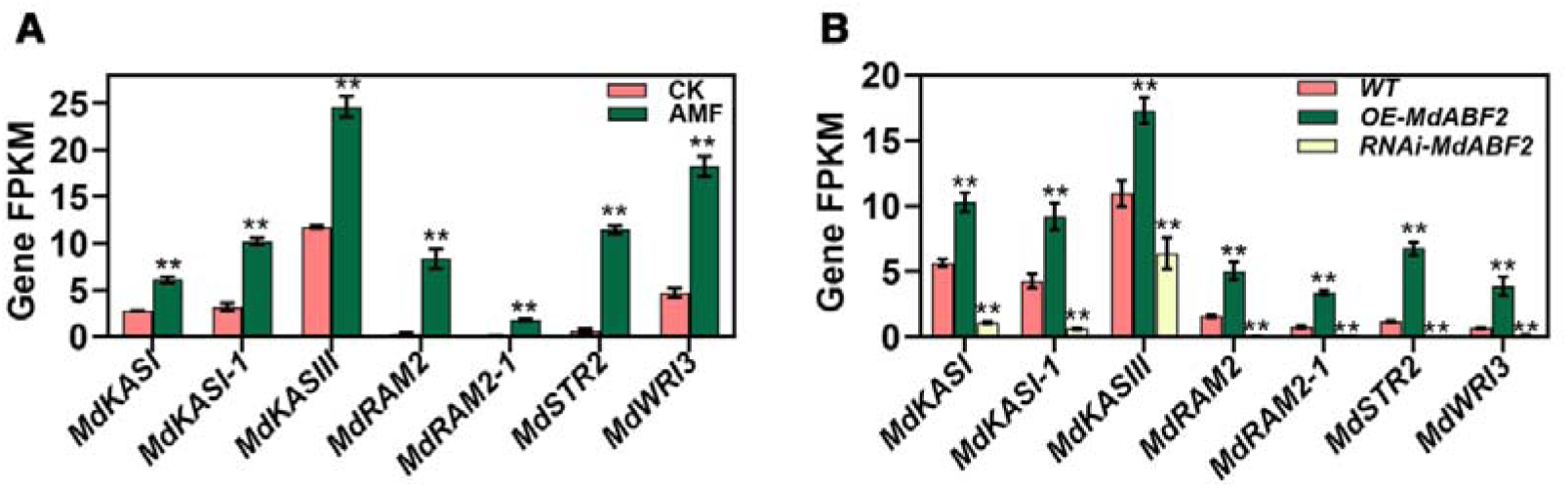
FPKM values for common fatty acid-associated genes that were differentially expressed at a significant level between two transcriptomes. **(A)** The FPKM values of differential gene expression levels related to fatty acids based on RNA-seq in the roots of 60-day-old uninoculated and inoculated M26 (*Malus pumila* Mill.) plants. **(B)** The FPKM values of differential gene expression levels related to fatty acids based on RNA-seq in WT and the *OE-MdABF2* apple (*Malus hupehensis* Rhed) transgenic hairy roots or the *RNAi-MdABF2* apple transgenic hairy roots. Expression-fold changes of DEGs (|log 2 FC| > 1, FDR < 0.05) represented significant difference. The bars represent the mean value ± SD (n = 3 independent biological replicates). Mixed samples from three apple seedlings carrying transgenic hairy roots were as one replicate.The asterisks indicate significant differences as assessed by one-way ANOVA (two-sided Student’s *t*-test; **P < 0.01).

**Supplemental Figure 10.**
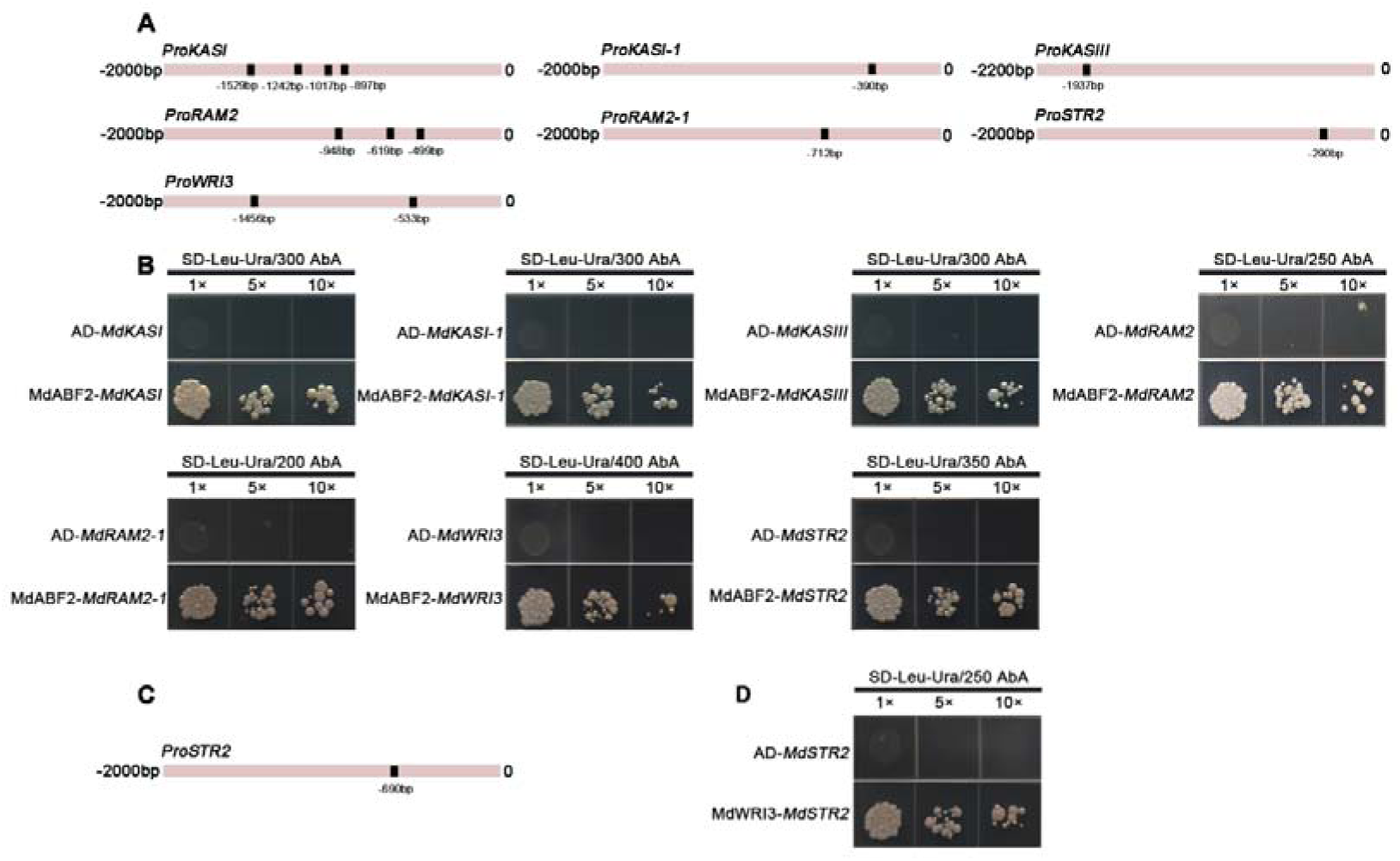
ABF2 interacts with the *MdKASI*, *MdKASI-1*, *MdKASIII*, *MdRAM2*, *MdRAM2-1*, *MdSTR2*, and *MdWRI3* promoters in a yeast one-hybrid assay. **(A)** Locations of predicted ABRE in the *MdKASI*, *MdKASI-1*, *MdKASIII*, *MdRAM2*, *MdRAM2-1*, *MdSTR2*, and *MdWRI3* promoters. **(B)** The Y1H assays demonstrated that MdABF2 directly binds to the *MdKASI*, *MdKASI-1*, *MdKASIII*, *MdRAM2*, *MdRAM2-1*, *MdSTR2*, and *MdWRI3* promoters. **(C)** Locations of predicted AW-box in the *MdSTR2* promoters. **(B)** The Y1H assays demonstrated that MdWRI3 directly binds to the *MdSTR2* promoters. *pGADT7* (AD) was used as a negative control.

**Supplemental Figure 11.**
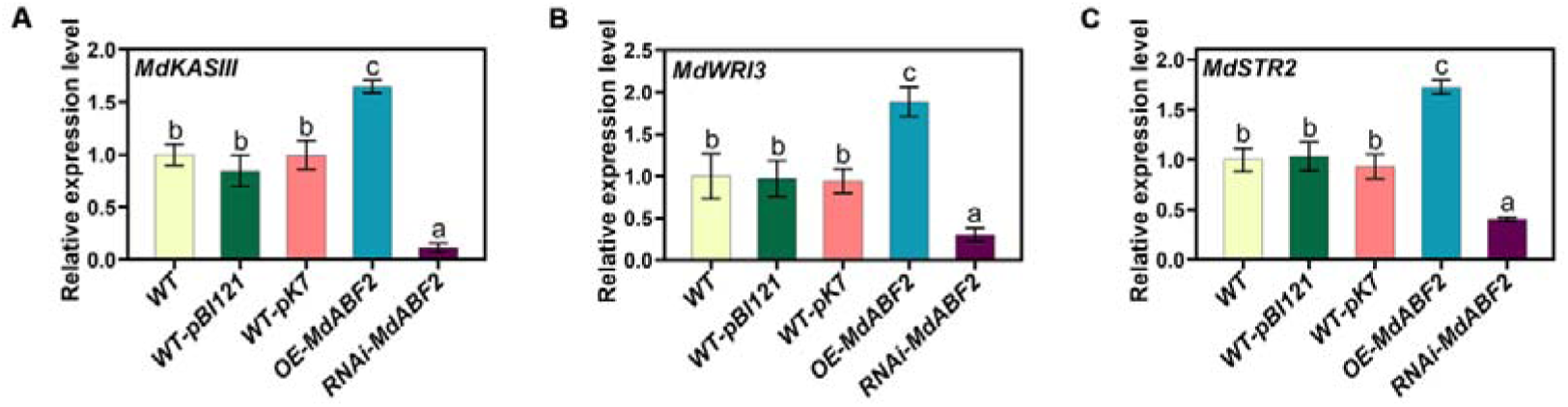
The relative gene expression levels in the roots of 60-day-old apple (*Malus hupehensis* Rhed) transgenic seedlings with *R*. *irregularis* infection of the transgenic roots. **(A–C)** The relative expression level of *MdKASIII* **(A),** *MdWRI3* **(B),** and *MdSTR2* **(C)** based on qPCR in WT, WT-pBI121, WT-pK7, *OE-MdABF2*, and *RNAi-MdABF2* apple transgenic hairy roots. Relative expression levels for each gene were obtained via the ddCT method, with its expression in WT set as ‘1’. The bars represent the mean value ± SD (n = 3 independent biological replicates). Mixed samples from three apple seedlings carrying transgenic hairy roots were as one replicate. Different letters indicate significant difference (analysis of variance [ANOVA]), Duncan’s multiple range test; *P* < 0.05).

**Supplemental Figure 12.**
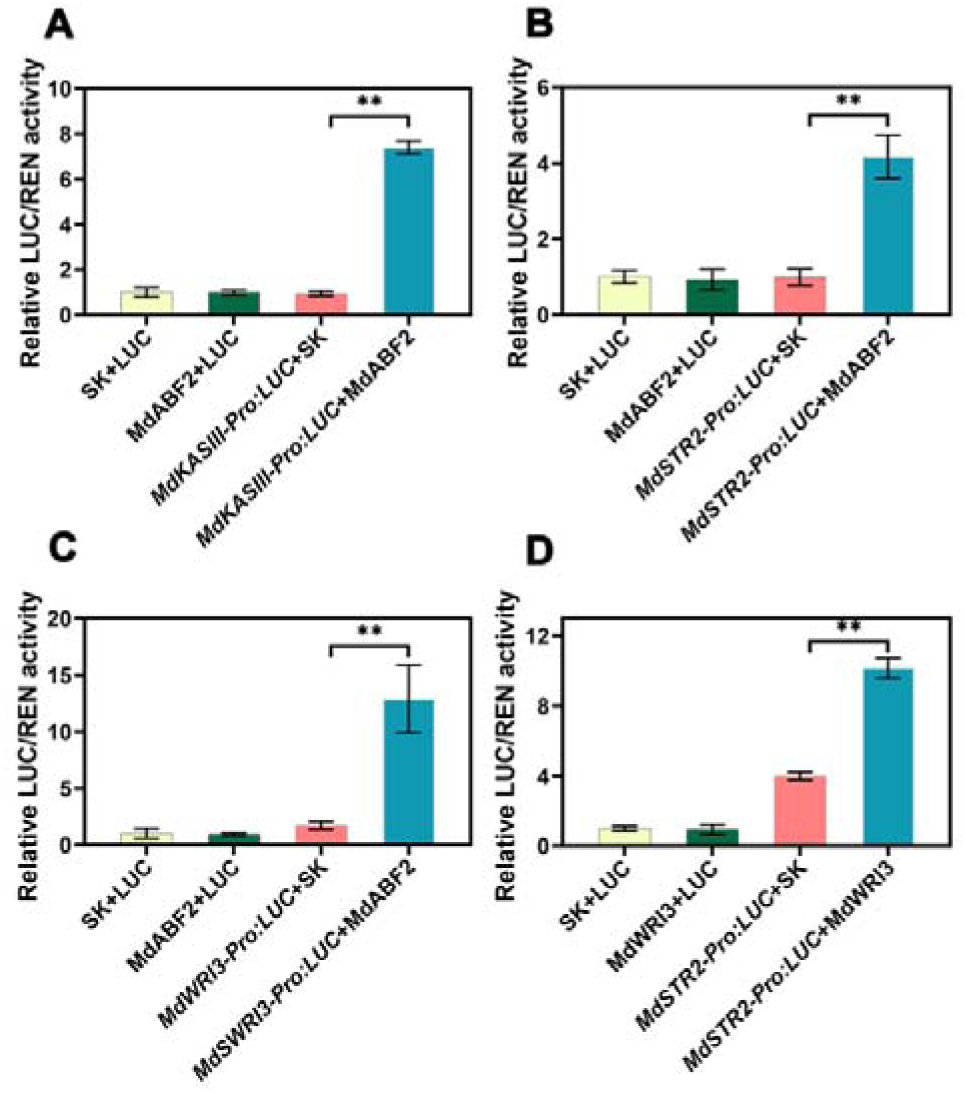
Relative LUC/REN activity of reporter and effector vectors for the LUC assays via transient expression assays in tobacco. **(A)** Relative LUC/REN activity of co-expressing MdABF2 and *MdKASIII-Pro:LUC*. **(B)** Relative LUC/REN activity of co-expressing MdABF2 and *MdSTR2-Pro:LUC*. **(C)** Relative LUC/REN activity of co-expressing MdABF2 and *MdWRI3-Pro:LUC*. **(D)** Relative LUC/REN activity of co-expressing MdWRI3 and *MdSTR2-Pro:LUC*. Relative LUC/REN activity, with its activity in SK+LUC set as ‘1’. The bars represent the mean value ± SD (n = 3 independent biological replicates). Mixed samples from three transgenic tobacco lines were as one replicate. The asterisks indicate significant differences as assessed by one-way ANOVA. (two-sided Student’s *t*-test; **P < 0.01)

**Supplemental Figure 13.**
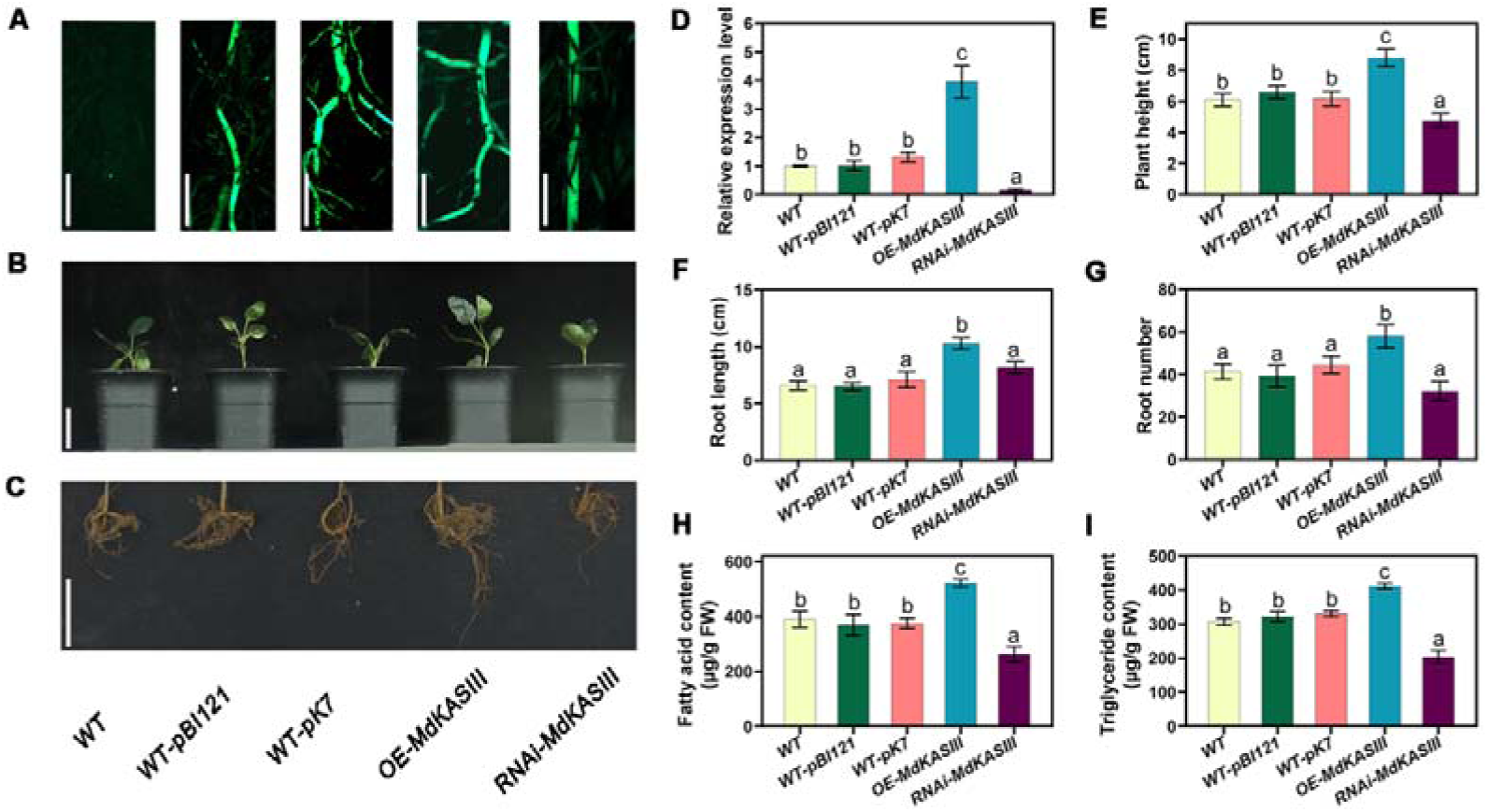
The effects of changing the expression of *MdKASIII* in the roots of 60-day-old apple (*Malus hupehensis* Rhed) transgenic hairy roots not inoculated with AMF. **(A)** Images of the transgenic root systems with green fluorescent protein. Scale bars, 1 mm. **(B)** Above-ground growth phenotypes of apple transgenic hairy roots. Scale bars, 5 cm. **(C)** Root growth phenotypes of apple transgenic hairy roots. Scale bars, 5 cm. **(D)** The mRNA relative expression levels of *MdKASIII* in the transgenic roots compared to the WT control (set as ‘1’). **(E)** The plant height of apple transgenic hairy roots. **(F)** The root length of apple transgenic hairy roots. **(G)** The root number of apple transgenic hairy roots. **(H)** The fatty acid content in the roots of apple transgenic hairy roots. **(I)** The triglyceride content in the roots of apple transgenic hairy roots. FW, fresh weight; WT, wild type; WT-pBI121, apple seedlings transformed with an empty overexpression vector containing the GFP tag (plasmid Binary Vector 121); WT-pK7, apple seedlings transformed with an empty RNA interference vector containing the GFP tag (pK7GWIWG2); *OE-MdKASIII*, *MdKASIII*-overexpressing root lines; *RNAi-MdKASIII*, *MdKASIII*-RNA interference root lines. (**D**, **H**, and **I)** The bars represent the mean value ± SD (n = 3 independent biological replicates). Mixed samples from three apple seedlings carrying transgenic hairy roots were as one replicate. (**E**, **F**, and **G)** The bars represent the mean value ± SD (n = 9 independent biological replicates). (**D**, **E**, **F**, **G**, **H**, and **I)** Different letters indicate significant difference (analysis of variance [ANOVA]), Duncan’s multiple range test; *P* < 0.05).

**Supplemental Figure 14.**
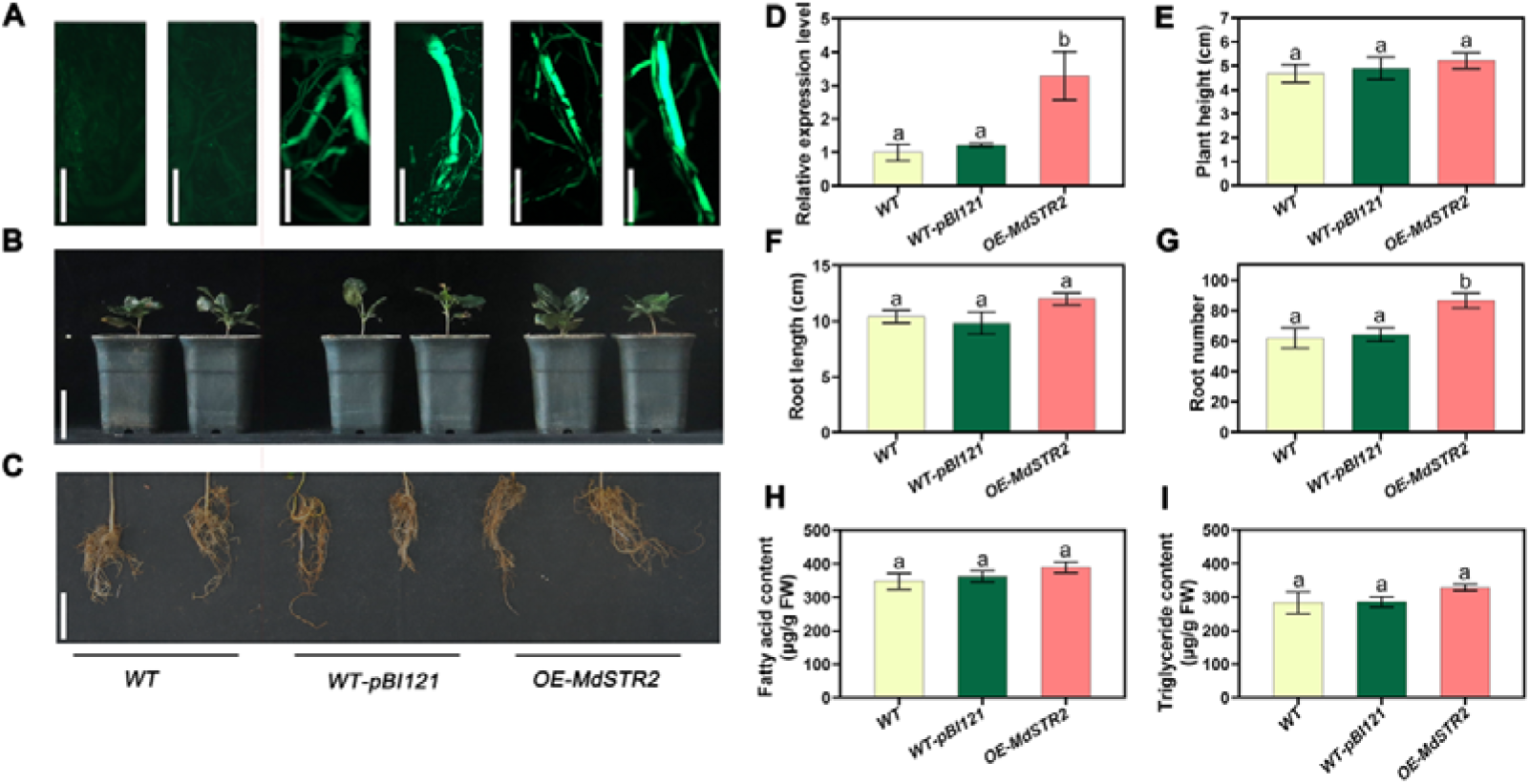
The effects of changing the expression of *MdSTR2* in the roots of 60-day-old apple (*Malus hupehensis* Rhed) seedlings not inoculated with AMF. **(A)** Images of the transgenic root systems of the apple seedlings with green fluorescent protein. Scale bars, 1 mm. **(B)** Above-ground growth phenotypes of apple transgenic hairy roots. Scale bars, 5 cm. **(C)** Root growth phenotypes of apple transgenic hairy roots. Scale bars, 5 cm. **(D)** The mRNA relative expression levels of *MdSTR2* in the transgenic roots of the apple seedlings compared to the WT control (set as ‘1’). **(E)** Plant height of apple transgenic hairy roots. **(F)** Root length of apple transgenic hairy roots. **(G)** Root number of apple transgenic hairy roots. **(H)** The fatty acid content of transgenic roots in the apple seedlings. **(I)** The triglyceride content of transgenic roots in the apple seedlings. FW, fresh weight; WT, wild type; WT-pBI121, apple seedlings transformed with an overexpressed empty vector containing the GFP tag (plasmid Binary Vector 121); OE-*MdSTR2*, *MdSTR2*-overexpressing root lines. (**D**, **H**, and **I)** The bars represent the mean value ± SD (n = 3 independent biological replicates). Mixed samples from three apple seedlings carrying transgenic hairy roots were as one replicate. (**E**, **F**, and **G)** The bars represent the mean value ± SD (n = 9 independent biological replicates). (**D**, **E**, **F**, **G**, **H**, and **I)** Different letters indicate significant difference (analysis of variance [ANOVA]), Duncan’s multiple range test; *P* < 0.05).

